# Cannabidiol reduces the latency for the behavioral effect of escitalopram in chronically stressed male mice: involvement of NAPE-PLD expressed in parvalbumin-positive interneurons and the prefrontal cortex

**DOI:** 10.1101/2021.04.23.441143

**Authors:** Franciele F. Scarante, Vinícius D. Lopes, Eduardo J. Fusse, Maria A. Vicente, Gabriel H. D. de Abreu, Viviani Nardini, Carlos A. Sorgi, Pedro H. C. Lirio, Feng Guo, Hui-Chen Lu, Jaime E. C. Hallak, Jose A Crippa, Lucia H. Faccioli, Katiuchia U. Sales, Francisco S. Guimarães, Ken Mackie, Alline C. Campos

## Abstract

Antidepressant drugs are the first-line treatment for chronic stress-related psychiatric disorders such as major depressive disorder, anxiety disorders, and post-traumatic stress disorder. However, their delayed-onset of therapeutic action, frequently occurring side effects, and incomplete clinical efficacy impose significant challenges for clinicians and patients’ adherence to treatment. Cannabidiol (CBD) is a major non-psychotomimetic phytocannabinoid with a wide range of potential clinical applications such as either a standalone drug or as an add-on treatment. In our study, we found that in chronically stressed male mice, CBD (30 mg/kg) rapidly induced behavioral improvement within 7 days, which was quicker than the high dose of escitalopram (ESC, 14 days). Additionally, repeated administration of a low and initially ineffective dose of CBD (7.5 mg/kg) potentiated the anti-stress effects of ESC (10 mg/kg) in mice subjected to 10 or 21 days of chronic unpredictable stress (CUS). Furthermore, our results suggested the involvement of N-acyl phosphatidylethanolamine phospholipase (NAPE-PLD) located in the prefrontal cortex (PFC) in the anti-stress effects of the 7-day treatment with ESC + CBD. This combination restored CUS-induced decreased expression of NAPE-PLD in the PFC. The behavioral effects of ESC + CBD were not observed in either constitutive NAPE-PLD knockout (KO) mice or mice with a CRISPR/Cas9-induced deletion of NAPE-PLD in the PFC. ESC + CBD treatment facilitated NAPE-PLD expression in parvalbumin (PV) interneurons in the PFC. As a conclusion, we suggest that CBD might be useful as an add-on therapy to optimize the action of (SSRI-)antidepressants, possibly by restoring the inhibitory/excitatory balance of the PFC via NAPE-PLD-mediated signaling.

**Highlights:**

- CBD (7.5 mg/kg) reduces the latency for anti-stress effects of escitalopram (ESC)
- ESC + CBD increases neuroplasticity in the prefrontal cortex (PFC)
- ESC + CBD reverses stress-induced loss of NAPE-PLD in PFC-Parvalbumin (PV)+ interneurons.
- NAPE-PLD in the PFC participates in the anti-stress effects of ESC + CBD.

**Graphical abstract:** Cannabidiol (CBD) enhances the antidepressant-like effects of escitalopram (ESC) in chronically stressed mice. While an effective dose of CBD (30mg/kg) alone rapidly improved stress-related behaviors when compared to a high dose of ESC (20mg/kg), ESC+CBD combination in sub-effective doses potentiated anti-stress responses and restored prefrontal cortex (PFC) function. These effects depended on N-acyl phosphatidylethanolamine phospholipase D (NAPE-PLD) activity within PFC parvalbumin interneurons, highlighting NAPE-PLD-mediated signaling as a key mechanism by which CBD may optimize antidepressant efficacy and reestablish inhibitory/excitatory balance in the PFC.

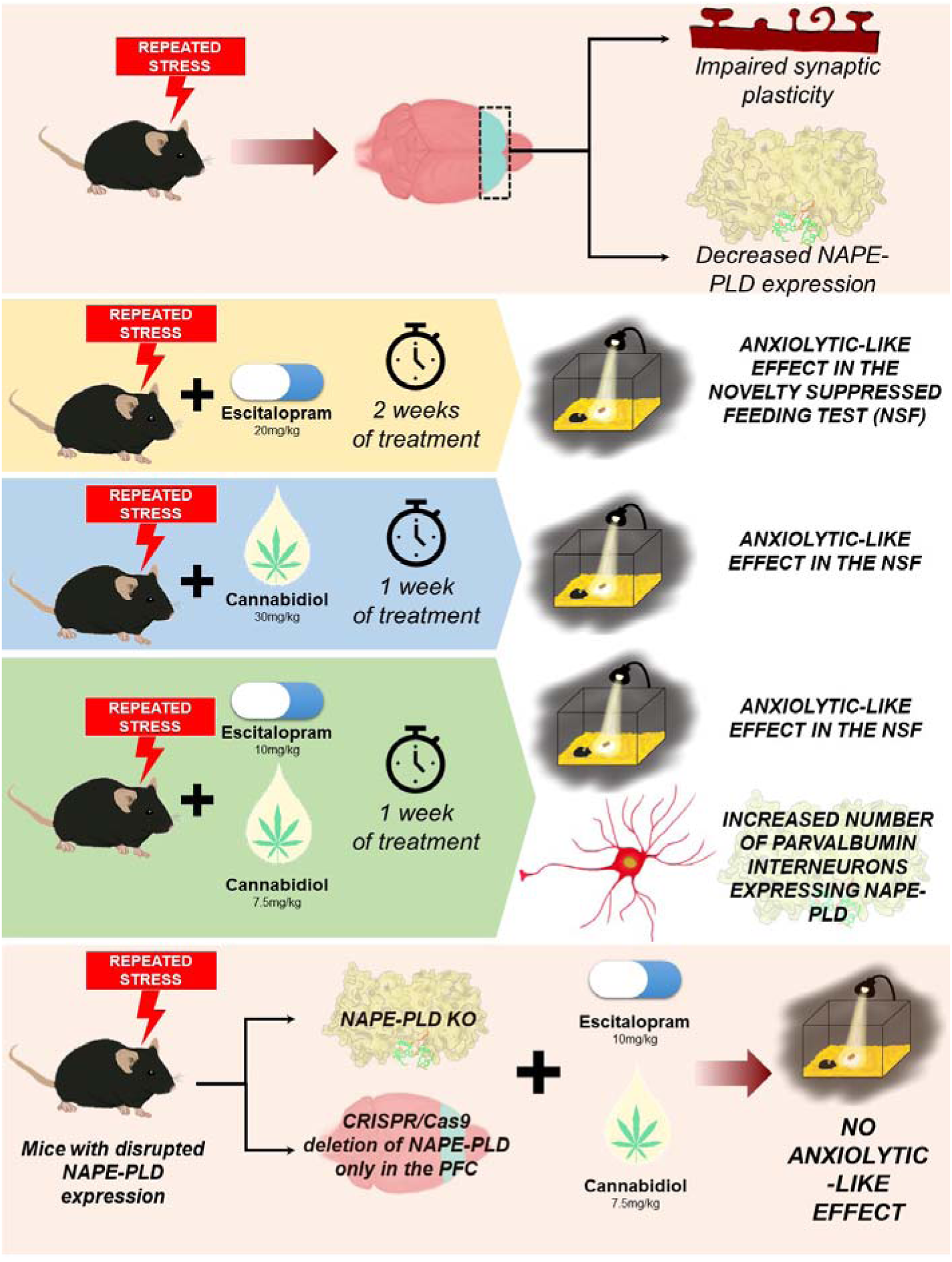

**Chemical compounds used in this article:** Cannabidiol (PubChem CID: 644019); Escitalopram oxalate (PubChem CID: 146571); URB597 (PubChem CID: 1383884); Ketamine hydrochloride (PubChem CID: 15851); xylazine hydrochloride (PubChem CID: 68554); 2,2,2-Tribromoethanol (PubChem CID: 6400); Flunixin meglumine (PubChem CID: 39212); Lidocaine hydrochloride (PubChem CID: 6314); Amoxicillin (PubChem CID: 33613).

## 1. Introduction

Selective serotonin reuptake inhibitors (SSRIs) are the most widely prescribed class of antidepressants (ADs) for the treatment of psychiatric disorders such as major depressive (MDD), anxiety and posttraumatic stress disorders (Caraci et al., 2018). However, their delayed onset of action, and association with significant and dose dependent side-effects and incomplete efficacy challenge patient adherence to SSRI therapy (Johnson et al., 2022). Furthermore, about 30% of patients are considered refractory or treatment-resistant to SSRIs (Mrazek et al., 2014), a situation often associated with significant burdens, including cognitive impairment, reduced quality of life, and increased suicide rates (Mrazek et al., 2014, Thase, 2011)

In studies using rodent models, SSRI treatment produces antidepressant- and anxiolytic-like effects, counteracting the behavioral and neuroplastic consequences of stress protocols (Satarelli et al., 2003). However, these effects generally emerge only after chronic treatment (21–28 days), reflecting the delayed onset of SSRI action observed in patients (Stassen et al., 1997). The time required for the induction of behavioral anti-stress effects by pharmacological treatments may be influenced by the specific brain region and neuroplastic mechanism targeted (Bambico et al., 2012). Traditional, antidepressants like SSRIs may rely on the induction of neuroplasticity in different brain areas such as the hippocampus. A classic example of antidepressant-induced hippocampal neuroplasticity is the process of adult neurogenesis, a considerably slower neuroplastic process compared to the synaptogenesis (Bambico et al., 2012). To date, only a few pharmacological strategies have been investigated to potentiate the effectiveness of antidepressants, such as the SSRI escitalopram, with the aim of reducing the 4–8-week latency of therapeutic effects without significantly increasing the incidence of side effects.

The phytocannabinoid cannabidiol (CBD), the major non-psychotomimetic component of the *Cannabis sativa* plant, also produces anti-stress effects and facilitates hippocampal neuroplasticity in rodents after 14 days (Campos et al., 2013, Fogaça et al., 2018). In addition, CBD has been shown to induce a rapid antidepressant-like effect in rodents submitted to behavioral tests sensitive to acute treatment of antidepressants, such as the forced swimming test (Linge et al., 2016, Sales et al., 2019). Although the molecular mechanisms that account for the behavioral and neuroplastic effects of CBD are currently under debate, they might involve the facilitation of anandamide (AEA) mediated neuromodulation, including the putative facilitation of its synthesis (via NAPE-PLD) and/or inhibition of its hydrolysis (FAAH) (Bisogno et al., 2001, Campos et al., 2013, Leishman et al., 2018). Interestingly, SSRIs have also been shown to modulate the function of the endocannabinoid system (Gunduz-Cinar et al., 2016, Smaga et al., 2014).

In the present study, we propose that CBD could be an effective add-on therapy to potentiate the behavioral and neuroplastic actions of SSRIs, significantly decreasing their slow onset of anti-stress action in male mice. Here, we tried to address the following questions: (a) Does CBD induce a faster behavioral effect compared to the SSRI escitalopram in stressed mice? (b) Can a sub-effective dose of CBD accelerate the behavioral and neuroplastic effects of escitalopram in stressed mice? (c) Can CBD help a SSRI to induce early neuroplastic effects in the PFC? (d) Which specific cell types are involved in this putative neurobiological mechanism?

## 2. Methods

### 2.1. Animals

The experiments involving animals described here were conducted following the current Brazilian guidelines for laboratory animals use (federal law number n° 11.794/08) and in compliance with the ARRIVE guidelines (Sert et al., 2020). All protocols were approved by the Ribeirão Preto Medical School Ethics Committee (Nb: 032/2015-1 and 047/2019) and the Bloomington Institutional Animal Care and Use Committee of Indiana University reviewed and approved the procedures used for Experiment 4 (SOP#52, 21-033). For experiments 1-3 and 5, C57Bl6 (2-3 months old; 24-30g) and Swiss male mice (6 months old-retired breeders) from the local animal facility of the University of São Paulo (Campus Ribeirão Preto) or from the animal facility from the Multidisciplinary Center for Biological Research (CEMIB; University of Campinas, Campinas – São Paulo, Brazil) were transferred to the animal care unit of the Pharmacology Department at least one month before the beginning of the experiments. For Experiment 4, WT and NAPE-PLD KO mice (generated by R. D. Palmiter and S. Luquet-Leishman et al., 2016) and WT mice (Jackson Labs) were of the same strain to which the KO mice were backcrossed into, the C57BL/6J strain. While the NAPE-PLD KO colony is maintained as homozygous crosses, once a year they were backcrossed to C57BL/6J (Jackson) mice to minimize genetic drift, and two non-sibling heterozygotes from this cross were bred to yield homozygous KOs. Animals were maintained in a temperature-controlled room (24°C) with water and food available *ad libitum* and under a 12h/12h light/dark cycle (lights on at 6h30). After arrival, they were housed in cages in groups of 3 (for social defeat stress protocols) or 4-5 (for chronic unpredictable stress protocols). Mice were assigned into experimental groups using the block randomization method.

### 2.2. Drugs

For the treatment schedule, we used escitalopram oxalate (ESC-Pratti-Donaduzzi; 10 and 20 mg/kg) (Ribeiro et al., 2023), Cannabidiol (CBD-99.9% purity-BSPG; 7.5 and 30 mg/kg) (Campos et al., 2013) and URB597 (Cayman; 0.1 mg/kg) (Fusse et al., 2024). Treatments were administered intraperitoneally (i.p.) daily for 7 or 14 days. Drugs were prepared daily (in 2% tween 80 + 98% saline solution) under sterile conditions at least 1h before the treatments and administered at 10mL/kg.

For the stereotaxic injections, mice were anesthetized with a single injection of 2,2,2-tribromoethanol (Sigma-Aldrich; 250mg/kg-i.p.) and received an additional local (cranial. Subcutaneous-s.c.) injection of lidocaine hydrochloride (Lidostesin®, Harvey) as well as an injection of Flunixin meglumine (Banamine®, Schering-Plough, 0,25mg/kg via s.c.). The post-surgical care included a 7-day treatment with amoxicillin (CIMED; 50mg/kg) orally administered via drinking water (0,25mg/ml).

Before perfusion and euthanasia, mice received a single injection (i.p., 10ml/kg of body weight) of a solution containing an overdose of ketamine hydrochloride (Syntex; 375mg/kg) and xylazine hydrochloride (Syntex; 25mg/kg).

### 2.3. Chronic unpredictable stress (CUS)

The model was adapted from Willner et al. (1992) as described in Campos et al. (2013) and Fogaça et al. (2019). Briefly, the animals were submitted daily to one of the stressors listed in Table 1 (distributed randomly). The stress procedure was conducted for 7, 10, 14 or 21 days, depending on the protocol. The control (non-stressed) mice were maintained at the animal care unit during all the experimental protocol, except during the behavioral tests.

**Table 1.** List of stressors randomly distributed in the CUS protocol. *In the 10-day CUS protocol those were the stressors that were applied more than once.

| Stressor | Duration |
| --- | --- |
| Forced swimming* | 15 minutes |
| Restraint stress* | 2 hours |
| Intermittent dark/light cycle | 20-24 hours |
| Inverted dark/light cycle | 20-24 hours |
| Wet bedding* | 20-24 hours |
| Tilted cage | Overnight |
| Food deprivation | 20-24 hour |

### 2.4. Social defeat stress (SDS)

The protocol used in our study was adapted from Golden et al. (2011) as described in Fusse et al., (2024). Before the stress procedure, C57Bl6 mice were housed in three animals per cage for at least one month before the start of the experiments. During this period, the animals established their social hierarchy within the home cage. Following this acclimation period, we implemented the Social Defeat Stress (SDS) procedure, which aimed to disrupt the established social hierarchy and induce a socially defeated phenotype in the resident mice. The SDS procedure involved introducing a Swiss intruder mouse, which was older and larger, into the cage of the resident C57Bl6 mice. This intrusion occurred for 2 hours per day over ten consecutive days, with a different intruder used daily for each resident cage. The Swiss mice underwent training and screening to assess their aggressiveness. The encounters were recorded, and at the end of the exposure, the mice were checked for possible injuries. The SDS mice were handled daily, and their behavior was evaluated by a blinded experimenter, who assessed factors such as locomotor activity, quality of self-cleaning behaviors, and aggressiveness. The control, non-stressed group was also housed in groups of three per cage in the same animal facility and mice were manipulated daily.

### 2.5. Novelty Suppressed Feeding Test (NSFT)

The test was performed as previously described Fogaça et al., 2018. Briefly, it consisted of placing 20-24h food-deprived mice in a rectangular arena (40 x 40 x 30 cm – the novelty component) located in a dark room. In the center of this arena, a food pellet was placed on an illuminated platform. To measure the novelty-induced hypophagia, we assessed the time elapsed before the animals started to eat the food pellet, with a cut-off time of 600 seconds. As a control measure of appetite, the latency to feed and food consumption in the mouse’s home-cage was evaluated for 5 minutes, right after the NSF test.

### 2.6. Experimental design

#### Experiment 1

This experiment was designed to address if CBD can produce a faster behavioral effect in the NSF than escitalopram in stressed mice. Mice were assigned in the following experimental groups (Stress condition/Treatment): (1) Control/Vehicle (n=9); (2) Control/CBD 30mg/kg (n=10); (3) Control/Escitalopram 20mg/kg (n=10); (4) CUS/Vehicle (n=10); (2) CUS/CBD 30mg/kg (n=8); and (3) CUS/Escitalopram 20mg/kg (n=10). Two independent protocols were conducted: 7-day CUS +treatment or 14-day CUS + treatment. On the 8^th^ or 15^th^ day of the treatment regimen, the NSF was conducted. On the following day, mice were submitted to transcardiac perfusion under deep anesthesia (ketamine 375mg/kg + xylazine 25mg/kg), and their brains were collected and processed for immunohistochemistry.

#### Experiment 2

Aiming to determine if combining a sub-effective dose of CBD can accelerate the anti-stress effects of escitalopram, mice were divided into the following experimental groups (Stress condition/Treatment 1/Treatment 2): (1) Control/Vehicle + Vehicle (CUS model: n=14; SDS Model: n=13); (2) Stressed/Vehicle + Vehicle (CUS model: n=11; SDS Model: n=8); (3) Stressed/Escitalopram 10mg/kg + Vehicle (CUS model: n=11; SDS Model: n=10); (4) Stressed/Vehicle + CBD 7.5mg/kg (CUS model: n=10; SDS Model: n=9); (5) Stressed/Vehicle + URB597 0.1mg/kg (CUS model: n=10; SDS Model: n=9); (6) Stressed/Escitalopram 10mg/kg + CBD 7.5mg/kg (CUS model: n=12; SDS Model: n=10); (7) Stressed/Escitalopram 10mg/kg + URB597 0.1mg/kg (CUS model: n=11; SDS Model: n=10). Two independent stress protocols were conducted: the CUS model and the SDS model. Both stress protocols were conducted for 10 days, and the drug treatments were conducted during the last 7 days of the stress protocol. All groups were tested in the NSF 24h after the last day of stress and treatment. Right after the behavioral assessments, a group of mice (n=4-6/group) was euthanized, and the tissues were dissected and processed for Golgi staining or western blot. Another group of mice were transcardially (n=5-6/group) perfused and had their brains collected and processed for immunohistochemistry.

#### Experiment 3

In this experiment we tested if a sub-effective dose of CBD could act as an add-on treatment to facilitate the anti-stress effects of ESC in mice submitted to a 21-day protocol of stress, while receiving pharmacological interventions from Day14 to Day 21 of the CUS protocol. Mice were separated in the following experimental groups (Stress condition/Treatment 1/Treatment 2): (1) Control/Vehicle + Vehicle (n=11); (2) Stressed/Vehicle + Vehicle (n=9); (3) Stressed/Escitalopram 10mg/kg + Vehicle (n=8); (4) Stressed/Vehicle + CBD 7.5mg/kg (n=8); (5) Stressed/Escitalopram 10mg/kg + CBD 7.5mg/kg (n=9). The CUS protocol was conducted for 21 days and the treatment was administered concomitantly to the last 7 days of the stress protocol. All groups were tested in the NSF 24h after the last day of stress and treatment. On the following day, mice were transcardially perfused, and the brains were collected and processed for immunohistochemistry.

#### Experiment 4

WT and NAPE-PLD KO mice were designated in the following experimental groups (Genotype/Stress condition/Treatment 1/Treatment 2): (1) WT/Control/Vehicle + Vehicle (n=7); (2) WT/Stressed/Vehicle + Vehicle (n=7); (3) WT/Stressed/Escitalopram 10mg/kg + Vehicle (n=7); (4) WT/Stressed/Escitalopram 10mg/kg + CBD 7.5mg/kg (n=7); (5) NAPE-PLD KO/Control/Vehicle + Vehicle (n=7); (6) NAPE-PLD KO/Stressed/Vehicle + Vehicle (n=8); (7) NAPE-PLD KO/Stressed/Escitalopram 10mg/kg + Vehicle (n=8); (8) NAPE-PLD KO/Stressed/Escitalopram 10mg/kg + CBD 7.5mg/kg (n=8). The CUS protocol was conducted for 10 days, and the treatments were administered during the last 7 days of the stress protocol. All groups were tested in the NSF 24h after the last day of stress and treatment.

#### Experiment 5

Animals received a bilateral intra-PFC injection of viral vectors delivering either a construct that induces the CRISPR-Cas9-mediated deletion of NAPE-PLD or DAGL-α (for the knock-down (KD) groups) or a scrambled sequence (for the wild-type – WT – groups) (n=15-16 for the NAPE-PLD KD experiment; n=9-10 for DAGL KD experiment). Fifteen days after the stereotaxic injection, mice were submitted to 10 days of CUS and 7 days of treatment with combinations of Vehicle or Escitalopram 10mg/kg with Vehicle or CBD 7.5mg/kg. Twenty-four hours after the last day of CUS and treatment, the behavioral response of mice was assessed in the NSF. For the control of the injection site, at the end of the stress and treatment protocols, the brains were sectioned (20µm), and the samples contained the PFC observed under the epifluorescence microscope (BX53, Olympus, Tokyo, Japan) to verify the position of the injection site according to the presence of tGFP.

### 2.7. Viral vectors

Lentiviral vectors were used to guide the CRISPR/Cas9 system and guide RNA sequence targeting the genes coding the NAPE-PLD enzyme or the DAGLα enzyme. For both targets, the vector consisted of a GeCKO2 lentiguide All in One system with coexpression of Cas9 and tGPF directed by the EF1α promoter and the expression of the guide RNA (gRNA) directed by the U6 promoter (GeCKO2 lentiguide-tGFP, U6-gRNA / EF1a-tGFP; NAPE-PLD Target ID: MM0000610807; NAPE-PLD Exon target: 3; DAGLa Target ID: MM0000628057; DAGLa Exon target: 10; Sigma-Aldrich). As a control, the CRISPR-lenti lentiviral vector non-directed-scramble sequence (control; Sigma-Aldrich) was used. The control plasmid co-expresses Cas9, GFP, and puromycin N-acetyltransferase directed by the EF1α promoter, while the U6 promoter directs the expression of the gRNA scrambled sequence.

### 2.8. Stereotaxic surgery for CRISPR-Cas-9 system injection into the prefrontal cortex

Eight-week-old mice were anesthetized with tribromoethanol (250mg/kg) and placed in a stereotaxic apparatus. We avoided the use of ketamine as our anesthetics due to its well characterized rapid and long lasting behavioral and neuroplastic effects in rodent models of chronic stress (Li et al., 2010). The coordinates used to reach the prefrontal cortex were (from Bregma, based on the Franklin and Paxinos, 2008): anteroposterior: (+) 1.5; mediolateral: (±) 0.4; dorsoventral: (-) 2.5. The injection of the previously described viral vectors was performed bilaterally (200μl per hemisphere) at a speed of 20nl/min using a Hamilton syringe coupled to an infusion pump. The needle was slowly removed from the site 2 minutes after the end of the infusion. Then, the animal was carefully removed from the apparatus. The infusion site was sutured with 4.0 nylon or silk thread. At the end of the surgery, each animal was placed in a post-surgical room for recovery. Mice were monitored for at least 2 hours and 1 hour after awakening from anesthesia. On the following 7 days after surgery, mice received daily treatment with amoxicillin in the drinking water (Marx et al., 2014).

### 2.9. Western blot

After euthanasia under deep anesthesia, the mice of experiment 2 had their PFC manually dissected on ice and harvested into 1.5mL plastic tubes. The samples were then homogenized in lysis buffer (Tris-HCl 25mM pH=7.5, NaCl 75mM, Na_2_H_2_O_7_P_2_ 2.5mM, Na_3_VO_4_ 1mM, Na_2_MoO_4_5mM, 0.5% Triton X-100) with protease inhibitor cocktail (*Protease Inhibitor Cocktail powder*, Cat. P2714, Components: AEBSF, Aprotinin, Bestatin, E-64, EDTA, Leupeptin; Sigma Aldrich, St. Loius, USA) using the bead-beater homogenizer (*TissueLyser LT Adapter*, Qiagen; 40Hz, 90 seconds). After the lysis, the samples were centrifuged (12000 rpm, 10 minutes, 4°C) the supernatant was collected for another round of centrifugation (12000 rpm, 10 minutes, 4°C). Bradford’s method was used to evaluate total protein concentration against a BSA standard curve and using a NanoDrop (absorbance reading at 450nm – *NanoDrop One Microvolume UV-Vis Spectrophotometer Thermo Scientific Waltham,* Massachusetts, USA). For protein separation, electrophoresis was conducted on a polyacrylamide gel with a gradient density (4-12% - NuPAGETM Invitrogen), with a final sample volume of 20uL and a protein concentration of 1g / L per well. After protein transfer onto a nitrocellulose membrane (Amersham Potran, Little Chalfont, United Kingdom), membranes were blocked in 10% non-fat milk (Bio-Rad) (dissolved in Tris-saline buffer + 0.5% of Tween20 -TBSt) for two hours, at room temperature. After blocking, membranes were rinsed quickly with TBSt to remove the excess of blocking solution and then incubated with the primary antibody at 4°C overnight under constant orbital agitation at the following dilutions: synaptophysin (∼33kDa; 1:1000; Rabbit; *Cell Signalling,* Cat. 5461S, Danvers, Massachusetts, USA), synaptotagmin (∼60kDa; 1:1000; Rabbit; *Cell Signalling,* Cat. 14558S, Danvers, Massachusetts, USA), PSD95 (∼95kDa; 1:1000; Rabbit; *Cell Signalling,* Cat. 3409S, Danvers, Massachusetts, USA) and α-tubulin (∼52kDa; 1:10000; Mouse; *Cell Signalling*, Cat. 3873S, Danvers, Massachusetts, USA). The secondary antibody was incubated for 2 hours (*Anti-rabbit*, *GE Healthcare*, 1:10000 or *Anti-mice*, *GE Healthcare*, 1:10000, Chicago, Illinois, USA). The bands were detected using chemiluminescence methods (ECLPrime®, Amersham, Little Chalfont, United Kingdom) and visualized using the ChemiDoc Imaging Systems (GE ImageQuant LAS, MA- United States). Intensities of specific bands were quantified using the Image Studio Lite (LI-COR, Nebraska, United States) and normalized to anti-α-tubulin protein levels.

### 2.10. Immunohistochemistry

#### Tissue processing

After anesthesia with ketamine and xylazine, mice from experiments 2 and 3 were transcardially perfused with 30mL of PBS 0.01M (pH=7.4-7.6) followed by 25mL of PFA 4% diluted in PB 0.2M. The brains were then dissected and maintained in PFA 4% for 24h before cryoprotection with 30% sucrose. brains were then frozen and maintained at −80°C until being sectioned in a cryostat. Six to eight sections (30µm) containing the vmPFC (From Bregma: −2.1 - −1.55mm- Paxinos & Franklin, 2001) were obtained and maintained in anti-freezing solution (ethyleneglycol 30% + glycerol 20%) at −20°C until the beginning of immunohistochemistry/fluorescence procedures.

#### Immunohistochemistry procedures

The immunostaining procedures were performed using the free-floating method. Six to eight slices containing different portions of the PFC were distributed in 24-well plates containing 1mL of TBS (50mM Tris, 150mM NaCl, pH 7.6). The protocol consisted of consecutive steps of: (1) Wash with TBS (1mL/well; 3x; 15 min, shaking); (2) Antigen retrieval with citrate buffer (1mL/well; pH 6.0) for 30 minutes at 70°C followed by cooling at room temperature (∼40minutes); (3) Wash with TBS (1mL/well; 3x; 15 min, shaking); (4) Blockade step in BSA 1% in TBS+0.25% Triton X-100 (1mL/well for 3-4h at room temperature; (5) Incubation with primary antibody solution diluted in TBS+ BSA 0.1% (250uL/well; 18h at 16°C, under constant shaking); (6) Wash with TBS+ (1mL/well; 4x; 15 min, shaking); (7) Incubation with secondary antibody solution diluted in TBS (2h at room temperature); (8) Wash with TBS+ (3x; 15 min, shaking); (9) Incubation with Hoescht (1:10000 from 2µg/mL stock solution, Invitrogen, Cat. H1399) diluted in TBS+ (15 min); (10) Mounting in glass slides using Fluormount-G (Electron Microscopy Sciences; Cat. 17984-25). The primary antibodies used in our study were anti-NAPE-PLD polyclonal antibody (1:400; Cayman Chemical Company, Cat. 10305) and anti-Parvalbumin mouse monoclonal antibody (1:500; Sigma-Aldrich, Cat. SAB4200545). The secondary antibodies used in our study were donkey anti-Rabbit IgG (H+L) Alexa Fluor 488 (1:1000; Thermo Fisher Scientific, Cat. A-21206) and goat anti-Mouse IgG (H+L) Alexa Fluor 647 (1:1000; Thermo Fisher Scientific, Cat. A-21235).

#### Image acquisition and analysis

Images were acquired with a confocal microscopy (Leica TS-SPE) and processed using the LasX software (v. 5.0.2; Leica Microsystem, Wetzlar, Germany). We obtained a relative measure of NAPE-PLD expression by puncta quantification of the NAPE-PLD staining using the Particle Analyser plug-in of the ImageJ software in the 3D image projections obtained coupling the z-stacks images after confocal scanning (Step size=0.3µm). We quantified the number of PV-positive cells and the percentage of PV cells expressing NAPE-PLD. The PV cells were counted when the whole cell body was visible in the area analyzed. We considered as a NAPE-PLD expressing PV cell, the ones with NAPE-PLD-puncta expression in the cell body. The results were normalized by the area and expressed as number of PV cells per mm² and as percentage of NAPE-PLD-expressing PV cells. We imaged and analyzed at least 6 slices/mouse corresponding to different portions of the vmPFC.

### 2.11. Golgi COX analysis

The Golgi impregnation method was performed using the FD Rapid GolgiStain Kit (FD NeuroTechnologies, INC) according to the manufacturer’s instructions. After the euthanasia under deep anesthesia, mice from experiment 3 had their brains removed from the skull and immersed in a potassium dichromate solution at room temperature for ten days. The tissues were then transferred to the washing solution and stored at room temperature for at least 72 hours before being sliced in a cryostat in 100μm sections placed on gelatinized slides. The slices were maintained hydrated until processing, which consisted of completing the reaction with mercury chloride and subsequent dehydration process in successive ethanol and xylene stages. The analysis was performed on pyramidal neurons of the mPFC layers II/III, selecting 10μm of the initial portion of tertiary dendrites. We quantified and classified the dendritic spines after scanning in the reflection mode of a confocal microscope (Leica TCS SPE; z step size: 0.44 µm; Reflection mode after excitation with 8% intensity of Laser 488). The analysis was performed using the LasX Leica Software, navigating through the images collected for each dendritic portion and quantifying the number and type of dendritic spines in the initial 10μm of the tertiary dendrites (Spiga er al., 2011).

### 2.12. Endocannabinoid quantification by LC -MS/MS

PFC samples were spiked with 10 pg of AEA-d8, homogenized in H2O/MeOH and purified as previously described (Bligh and Dyer, 1959). Following the extraction step, the samples were dried, suspended in 100 μl of MeOH, then applied into the LC– MS/MS. Mobile phases consisted in water or acetonitrile, both containing 0.1% formic acid. The gradient condition was 0 to 1 min, 5% in acetonitrile for 5 min, 70% in acetonitrile for 9.5 to 11 min, 98% in acetonitrile. A flow rate of 0.5 ml/min and Ascentis Express C8 (150 × 2.1 mm; 2.7 μm) column was employed. Mass spectrometry was performed in positive mode for MRMHR (high-resolution multiple reaction monitoring) analysis. Results are expressed as pmol per mg of tissue (de Oliveira et al., 2020)

### 2.13. Statistical analysis

All data obtained was tested for normality using the Kolmorov-Smirnov test and for homogeneity of variances using the Levene test. In the experiments involving the comparison of behavioral effects of CBD and escitalopram, we used the Two-way ANOVA (factor 1: stress; factor 2: treatment). In the experiments analyzing the effects of the combination of cannabinoids and escitalopram, the stress factor effect was assessed using the t-test for independent samples (comparing the vehicle-treated non-stressed with the vehicle-treated stressed group) (Fuse et al., 2022; Fernandes et al., 2021; Ribeiro et al., 2022). Then we used Two-way ANOVA to evaluate the effects of the factors: treatment 1 (vehicle or ESC), and treatment 2 (vehicle, CBD, or URB597) within the stress groups. Differences between groups were further analyzed using Duncan’s post-hoc test. For the experiments that involved the comparison of genotypes, we used p values equal to or less than 0.05 were considered significant. For every behavioral experiment, z scores were measured to detect outlier-subjects for each data point by taking the raw data, subtracting the mean, and dividing by the standard deviation. Data is represented as mean ± standard error of the mean (SEM). Mice were distributed in experimental groups using block randomization. The group size was calculated using G-Power software using beta=80% and alpha=0.05.

## 3. Results

### 3.1. Does cannabidiol produce a faster onset of behavioral effects than escitalopram in stressed mice?

To compare the onset of action of escitalopram (ESC) versus CBD in stressed mice, we assessed the behavioral effects of these compounds after 7 or 14 days of chronic unpredictable stress (CUS) exposure. In the 7-day CUS and treatment protocol, ESC (20mg/kg/day i.p.) reduced the latency to eat in the novelty-induced hypophagia in the control, but not in the stressed group (F_2,52_=6.442, p=0.003). On the other hand, CBD (30mg/kg/day i.p.), reduced latency to eat in the novel context after seven days of treatment (F_5,52_=4.596, p=0.002; One-way ANOVA followed by Duncan, Figure 1A). The One-way ANOVA results were reinforced by Log-rank (Mantel-Cox) test followed by Gehan-Breslow-Wilcoxon test (Mantel-Cox test: p=0.0002; Control Veh vs Control ESC: p=0.0318; CUS Veh vs CUS CBD: p=0.003; CUS Veh vs CUS ESC: p=0.0497) (Figure 1B). There was no stress or treatment effect on food consumption in the home-cage (Figure 1C).

**Figure 1.**
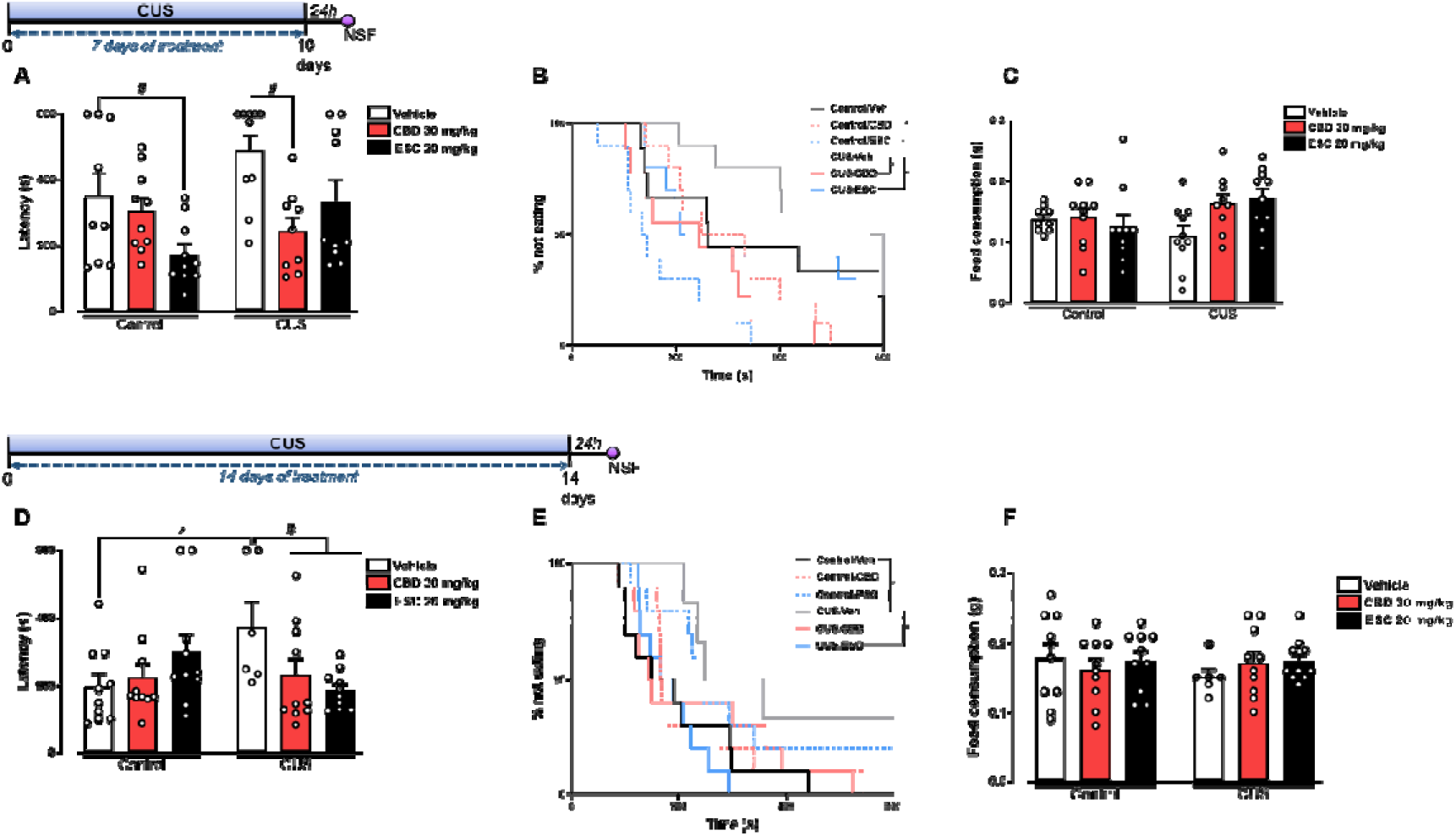
CBD produces a faster anti-stress effect than escitalopram. Effect of 7 (A-C) and 14 days (D-F) of treatment with escitalopram (20mg/kg) or CBD (30mg/kg) on the response of mice undergoing the chronic unpredictable stress (CUS) model in the latency to start feeding in the novelty suppressed feeding (NSF) test (A and D), on the percentage of mice not eating over the 600s of the test (B and E) and on the home-cage food consumption (C and F). (*) indicates p<0.05 compared to the control group; (#) indicates p<0.05 compared to the stressed group treated with vehicle (N=6-10, Two-way ANOVA; One-way ANOVA followed by Duncan; Mantel-Cox log-rank test followed by Gehan-Breslow-Wilcoxon test; t test for independent samples).

In the 14-day stress protocol, CBD’s effects on the stressed mice were maintained. After 14 days of daily treatment, ESC (20mg/Kg/Day i.p.) reduced latency to eat in the new environment. Neither of the treatments (ESC or CBD) affected the behavior of control animals (F_2,50_=4.778, p=0.013, Figure 1D). Figure 1E shows the survival curve analysis of the 14-day protocol. Therefore, CBD showed a more rapid onset of the anti-stress effect than escitalopram. No difference in food consumption in the home-cage was detected in 14 days (Figure 1F).

Since alterations in adult hippocampal neurogenesis have been observed after repeated treatment with CBD (Campos et al., 2013) and SSRIs (Ribeiro et al., 2022, 2023, Santarelli et al., 2003), we measured the number of cells positive for the neuroblast marker doublecortin (DCX) in the dentate gyrus (DG) after the 7- or 14-day protocols (Sup Figure 1). In non-stressed groups treated with the drug regimen of 14 days, CBD (30mg/kg), but not ESC (20mg/kg) increased the number of DCX+ cells (Supplementary figure 1 and Table1). However, no association (Supplemental Table 1) or correlations (data not shown) were found between the anti-stress effects and the number of DCX-positive cells in the DG.

### 3.2. Can a sub-effective dose of CBD shorten the onset of ESC action in stressed mice?

Next, we evaluated if a subeffective dose of CBD (7.5mg/kg/day i.p.) could act synergically to potentiate the anti-stress effects of a lower dose of ESC (10mg/kg/day i.p.), achieving efficacy in only 7 days. Mice were submitted to 10 days of CUS and treated with the combination of escitalopram and CBD for the last 7 days. Using this protocol, stressed mice treated with the combination ESC+CBD had a significantly shorter latency to feed in the new environment (F_5,60_=2.437, p=0.045, Figure 2A). This effect was further verified by Mantel-Cox log-rank test followed by Gehan-Breslow-Wilcoxon test (Mantel Cox: p=0.0008; Gehan-Breslow-Wilcoxon: CUS Veh/Veh vs CUS ESC+CBD: p=0.0405) (Figure 2B). No differences were detected in home-cage food consumption (Figure 2C). Since increased AEA levels have been associated with the behavioral effects of CBD in the CUS model [4, 5], we also sought to determine if URB597 (0.1mg/kg) in combination with escitalopram, would produce a similar add-on effect as CBD. However, we did not observe any effect (Figure 2A).

**Figure 2.**
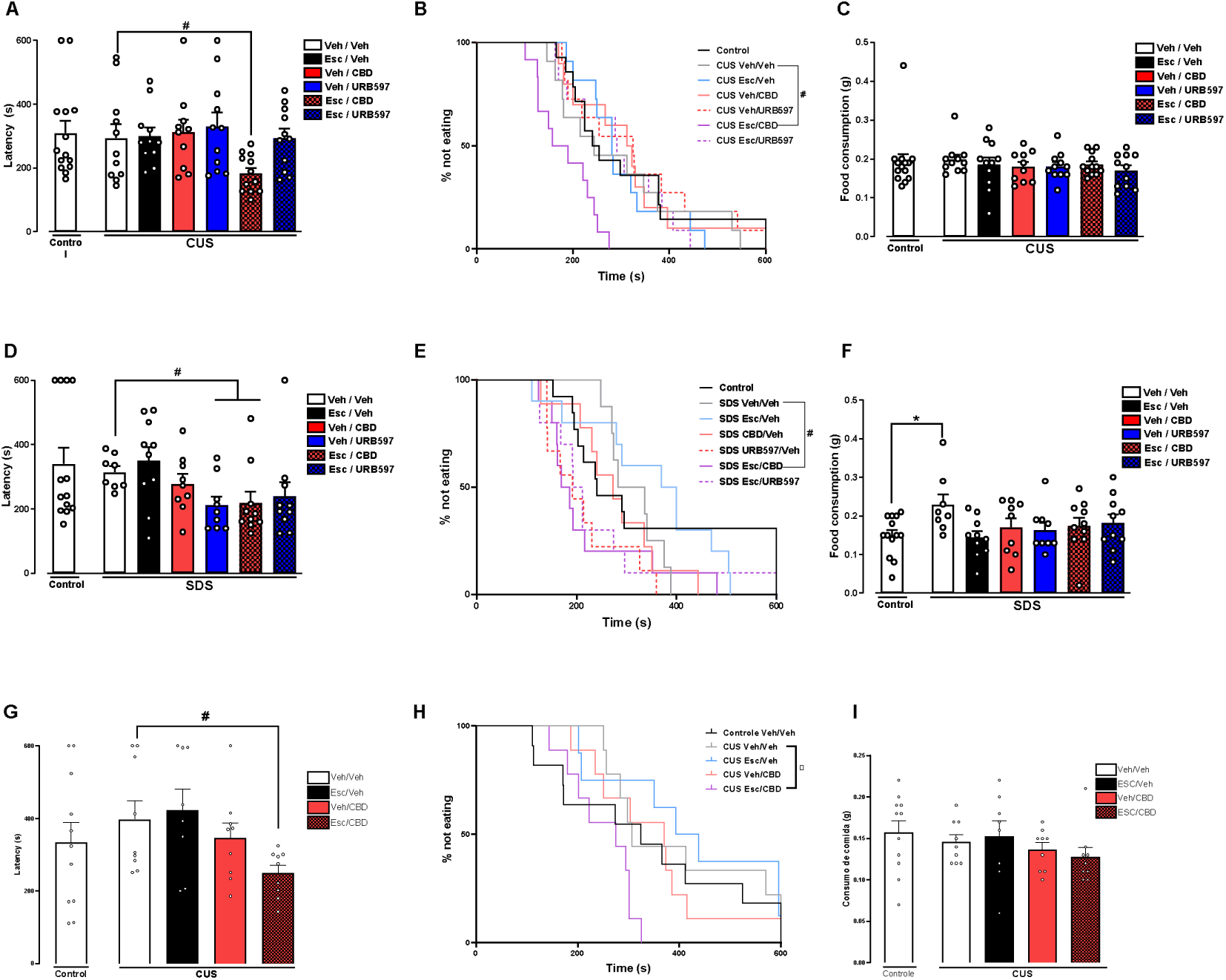
Sub-effective doses of CBD act as an add-on treatment, reducing the latency of onset of the anxiolytic-like effect of escitalopram in chronically stressed animals. Effect of the 7-day treatment with the different combinations of vehicle or escitalopram (10mg/kg) and vehicle, CBD (7.5mg/kg) or URB597 (0.1mg/kg) on the latency to start feeding in the NSF test (A, D and G), on the percentage of mice eating throughout the 600s of the test (B, E and H) and on the home-cage food consumption (C, F and I) in mice submitted to the CUS model for 10 (A-C) or 21 days (G-I) or exposed to the social defeat model (SDS) (D-F). Data represented as Mean ± SEM (A-H); (*) indicates p<0.05 compared to the control group; (#) indicates p<0.05 compared to the stressed group treated with vehicle (N=8-14, Two-way ANOVA; One-way ANOVA followed by Duncan; Mantel-Cox log-rank test followed by Gehan-Breslow-Wilcoxon test; t test for independent samples).

Thereafter, we tested if the add-on effects of CBD could be observed in mice submitted to another stress model, the SDS protocol. Again, the combination of ESC+CBD, but not ESC+URB, significantly decreased the latency to feed in the new environment (Figure 2D; F_5,50_=2,471, p=0.045, One-way ANOVA followed by Duncan). In socially stressed mice, URB597 also had a reduced latency in NSFT. Montel-Cox log-rank test followed by Gehan-Breslow-Wilcoxon test also indicated a difference between the SDS Veh/Veh group and the SDS Esc+CBD group (Mantel-Cox: p=0.0025; Gehan-Breslow-Wilcoxon: SDS Veh+Veh vs SDS ESC+CBD: p=0.0187) (Figure 2E). Moreover, the stressed vehicle-treated group consumed more food in the home-cage than non-stressed mice (Student’s t-test; t_19_=2.821, p=0.011). Still, there was an interaction between the treatments (F_2,49_=3.283, p=0.046) (Figure 2F). The SDS group treated with escitalopram alone presented a home-cage consumption significantly diminished compared to the SDS group that received only vehicle (One-way ANOVA followed by Duncan) (Figure 2D).

We also evaluated the effects of the treatments on stressed mice in the tail suspension test (TST) (Figure S2). Both CUS and SDS enhanced passive coping strategies in the TST, decreasing the latency to the first immobility episode (CUS: Student’s t-test; t_21_=2.652, p=0.015; SDS: Student’s t-test; t_15_=2.787, p=0.014). Interestingly, the SDS group treated with Escitalopram + CBD showed a significantly higher latency for the first immobility episode (F_5,44_=1.690, p<0.05; One-way ANOVA followed by Duncan) (Figure S2). This response is predictive of an antidepressant-like effect of the drug combination. Additionally, mice submitted to CUS and treated with vehicle presented an increase in overall immobility during the TST test (F_1,21_=7.863, p=0.011). In the SDS protocol, repeated measures ANOVA indicated an interaction between stress and time in the TST, indicating that the immobility response along the 6 minutes of the test varied depending on whether the mouse was or was not submitted to the homotypic stress model (F_5,75_=3.018, p=0.015). Concerning the immobility time, however, there were no significant differences between the stressed groups that received different treatment combinations neither in the CUS (repeated measures ANOVA; Treatment 1: F_1,54_=0.076, p=0.784; Treatment 2: F_2,54_=0.492, p=0.614; Interaction: F_2,54_=0.159, p=0.854) nor in the SDS protocol (repeated measures ANOVA; Treatment 1: F_1,42_=1.899, p=0.176; Treatment 2: F_2,42_=0.599, p=0.554; Interaction: F_2,42_=0.067, p=0.935) (Figure S2).

We also wondered, however, whether the combination with CBD would also impact the effects of escitalopram in reversing the effects of stress in mice exposed to 21 days of CUS, but treated with ESC+CBD in last 7 days of the stress protocol. Two-way ANOVA revealed that there was no effect of Treatment 1 (vehicle or escitalopram) (F_1,31_=1.138, p=0.294). On the other hand, there was a significant effect of Treatment 2 (Vehicle or CBD) (F_1,31_=6.074, p=0.019). No significant interaction between factors (F_1,31_=2.002, p=0.167). These results indicate that CBD as an add-on treatment alters the behavioral response of stressed mice. One-way ANOVA followed by Duncan post-hoc indicated that mice submitted to 21 days of CUS and treated for only 7 days with the combination of escitalopram and CBD display a significantly reduced latency to feed in a context of neophobia in comparison to the vehicle-treated stressed group (F_3,31_=3.090, p=0.041) (Figure 2G). Mantel-Cox log-rank test followed by Gehan-Breslow-Wilcoxon test also indicated a difference in the response of CUS Veh+Veh group and CUS EEC+CBD group in the NSF (Mantel-Cox: p=0.050; Gehan-Breslow-Wilcoxon: CUS Veh+Veh vs CUS Esc+CBD: p=0.043) (Figure 2H). Home-cage food consumption was not significantly altered by any of the factors (Stress: independent samples t test - t_16,093_=0.706, p=0.490; Two-Way ANOVA: Treatment 1: F_1,31_=0.007, p=0.936; Treatment 2: F_1,31_=1.963, p=0.171; Interaction: F_1,31_=0.436, p=0.514) (Figure 2I). Therefore, a 7-day treatment with the combination of escitalopram with a sub-effective dose of CBD was sufficient to promote behavioral effects in mice exposed to either 10 or 21 days of stress. Next, we sought to evaluate the involvement of prefrontal cortex (PFC) and hippocampus in the behavioral effects of CBD+ESC, by analyzing whether the observed behavioral effects were associated with changes in markers of synaptic plasticity. In the PFC, the ESC+ CBD SDS-group showed an increased expression of synaptophysin (Synaptophysin: F_2,18_=7.437, p=0.004). Interestingly, two-way ANOVA revealed a significant interaction between treatments concerning the expression of synaptotagmin in the PFC of stressed mice (F_2,18_=3.866, p=0.040), indicating that the treatment combination alters the expression of this presynaptic protein (Sup Figure 3A-D). We also evaluated the impact of the various treatments on the expression of the postsynaptic protein postsynaptic density 95 (PSD95) in the PFC. Neither the SDS-stress protocol nor the drug regimen treatments significantly altered the relative expression of this protein in PFC (Sup Figure 3H). The observed changes in synaptic proteins induced by the SDS protocol were not replicated by CUS.

In the hippocampus, the combination of escitalopram and CBD did not significantly alter the expression of synaptophysin but increased the relative expression of synaptotagmin in stressed mice (F_2,17_=4.242, p=0.032, Sup Figure 4C). However, the combination of escitalopram and URB597, did increase hippocampal synaptophysin (Sup Figure 4, F_2,17_=8.203, p=0.003), demonstrating a possible different neurobiological mechanism for CBD and UBB597 in modulating synaptic plasticity. Also, we did not observe changes in the markers of post-synaptic plasticity in the hippocampus (Sup. Figure 4E-H).

We also applied the Golgi staining technique to visualize and quantify the number and type of dendritic spines in pyramidal neurons of the layers II/III of the medial PFC (Sup. Figure 3E) and granule cells of the DG of the hippocampus (Sup. Figure 4E) as a measure of post-synaptic structural plasticity. Based on their morphology, the dendritic spines were classified as: mushroom, thin, filopodia, or stubby (Sup. Figure 3E-F; Sup. Figure 4E-F). The percentage of each subtype was quantified relative to the total number of dendritic spines (Sup Figure 3F; Sup. Figure 4F). The percentage of spines with thin morphology was increased in the PFC of the CUS group compared to the non-stressed animals (Sup Figure 3F; Student’s t-test; t_8_=2.097, p=0.020). CUS also decreased the total number of dendritic spines in tertiary dendrites of pyramidal neurons in the mPFC (t_5.219_=2.604, p=0.046, Sup Figure 3G). Again, there was no effect of treatment with the various drugs on spine morphology. Our data indicates that stress impairs structural postsynaptic plasticity in the prefrontal cortex (PFC). The combination of escitalopram and cannabidiol (CBD) may restore the expression of presynaptic proteins involved in synaptic transmission, thereby reestablishing PFC synaptic function. These findings suggest that the PFC could be a key neurobiological site underlying the behavioral effects of the ESC+CBD treatment and provide a basis for further molecular analyses in our study.

### 3.3. CBD+ESC restores stress-decreased NAPE-PLD levels in parvalbumin positive neurons within the prefrontal cortex

Several studies have speculated through associative data that CBD (30mg/kg) increases levels of AEA to promote anti-stress effects (Campos et al., 2013; Fogaça et al., 2018; Leweke et al., 2012). Curiously, AEA-mediated neuromodulation seems to be enhanced after repeated treatment with escitalopram (Smaga et al., 2014). Leishman et al. (2018) proposed that CBD enhances AEA via a NAPE-PLD-dependent mechanism, and Smaga and colleagues (2019) found that repeated treatment with escitalopram increases the expression of NAPE-PLD. Therefore, we next sought to investigate whether the neuroplastic alterations induced by the combination of escitalopram and CBD in the mPFC would be accompanied by changes in NAPE-PLD expression. We found that, in animals submitted to CUS (10 days-protocol) NAPE-PLD expression was reduced when compared to the control group (independent samples t test; 10-day CUS: t_7_=3.685, p=0.008; 21-day CUS:t_4_=2.166, p=0.096). Two-way ANOVA did not show a significant effect of Treatment 1 (F_1,14_=0.297, p=0.594), but there was a significant effect of Treatment 2 (F_1,14_=5.761, p=0.031) and a significant interaction between treatments (F_1,14_=5.784, p=0.031). One-way ANOVA followed by Duncan indicated that stressed mice treated with the combination of ESC+CBD showed an increased density of NAPE-PLD puncta in the PFC in comparison to the stress group treated with only escitalopram (Figure 3A-C).

**Figure 3.**
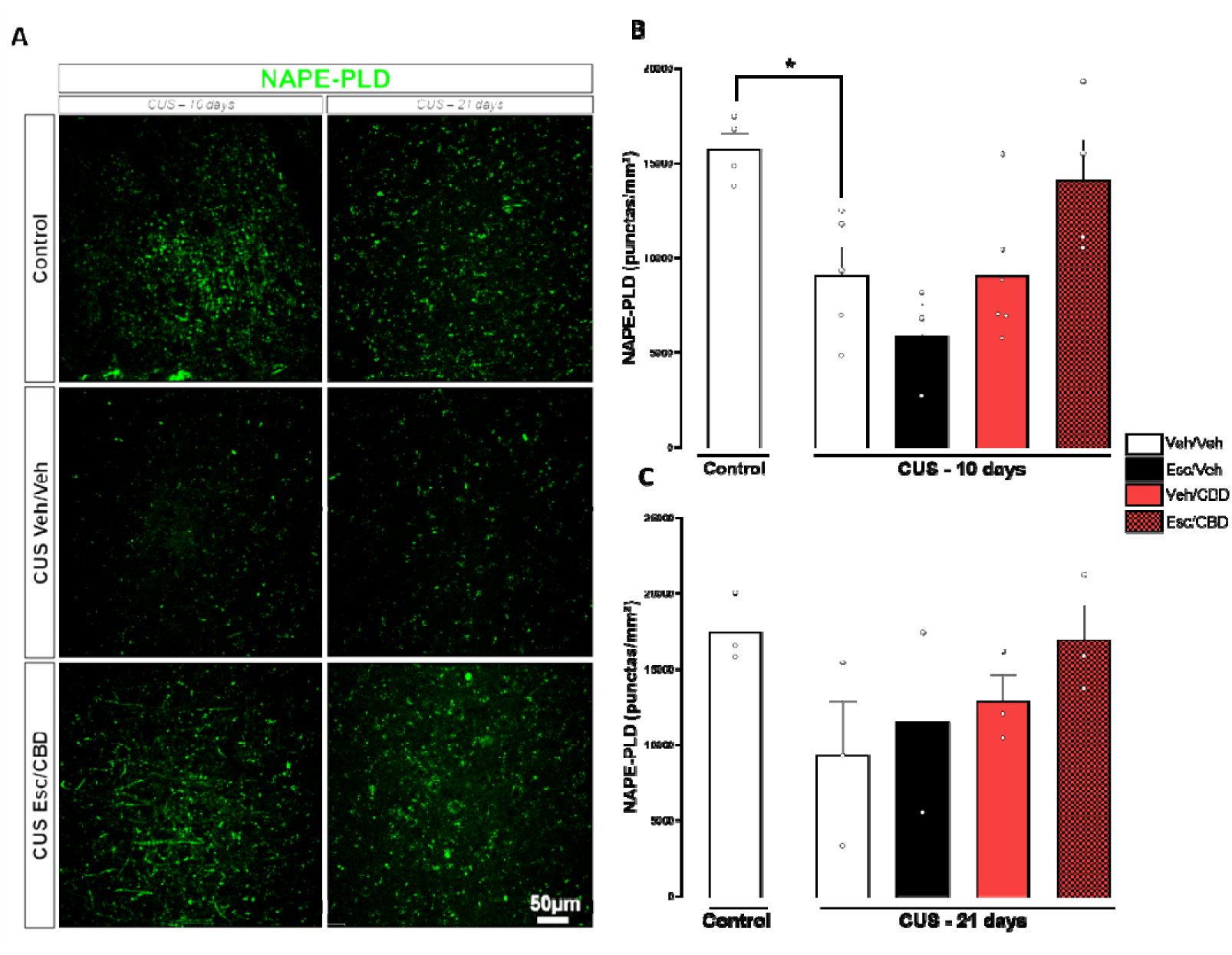
NAPE-PLD expression is reduced in the PFC of stressed mice. (A) Representative images of NAPE-PLD immunostaining; (B and C) Density of punctas of NAPE-PLD staining in the PFC of mice stressed for 10 (B) or 21 days (C) and treated for 7 days with the combinations of vehicle or escitalopram (10mg/kg) with vehicle or CBD (7.5mg/kg). Data represented as the Mean ± SEM; (*) indicates p<0.05 compared to the control group (t test for independent samples).

We next wondered if the pattern of expression of NAPE-PLD was altered in specific neuronal populations within the vmPFC. Our data suggested that treatment with the combination of escitalopram and CBD increased the percentage of PV-positive cells expressing NAPE-PLD around the soma in the vmPFC compared to the vehicle-treated group (One-way ANOVA followed by Duncan; F_3,7_=3.293, p=0.05- Figure 4A and C). We did not find significant changes in PV+ immunoreactivity in any of the conditions tested (Figure 4B), although qualitative analysis suggested a decrease in expression of this calcium binding protein in the dendrites of the PV+ neurons (figure 5A, upper panel), which seems to be partially recovered by ESC+CBD treatment (Figure 4A, lower panel). No differences concerning the pattern of NAPE-PLD expression in CamKII_α_-positive cells were detected (data not shown). Therefore, the treatment combination seems to increase the expression of NAPE-PLD in PV-positive interneurons in the vmPFC.

**Figure 4.**
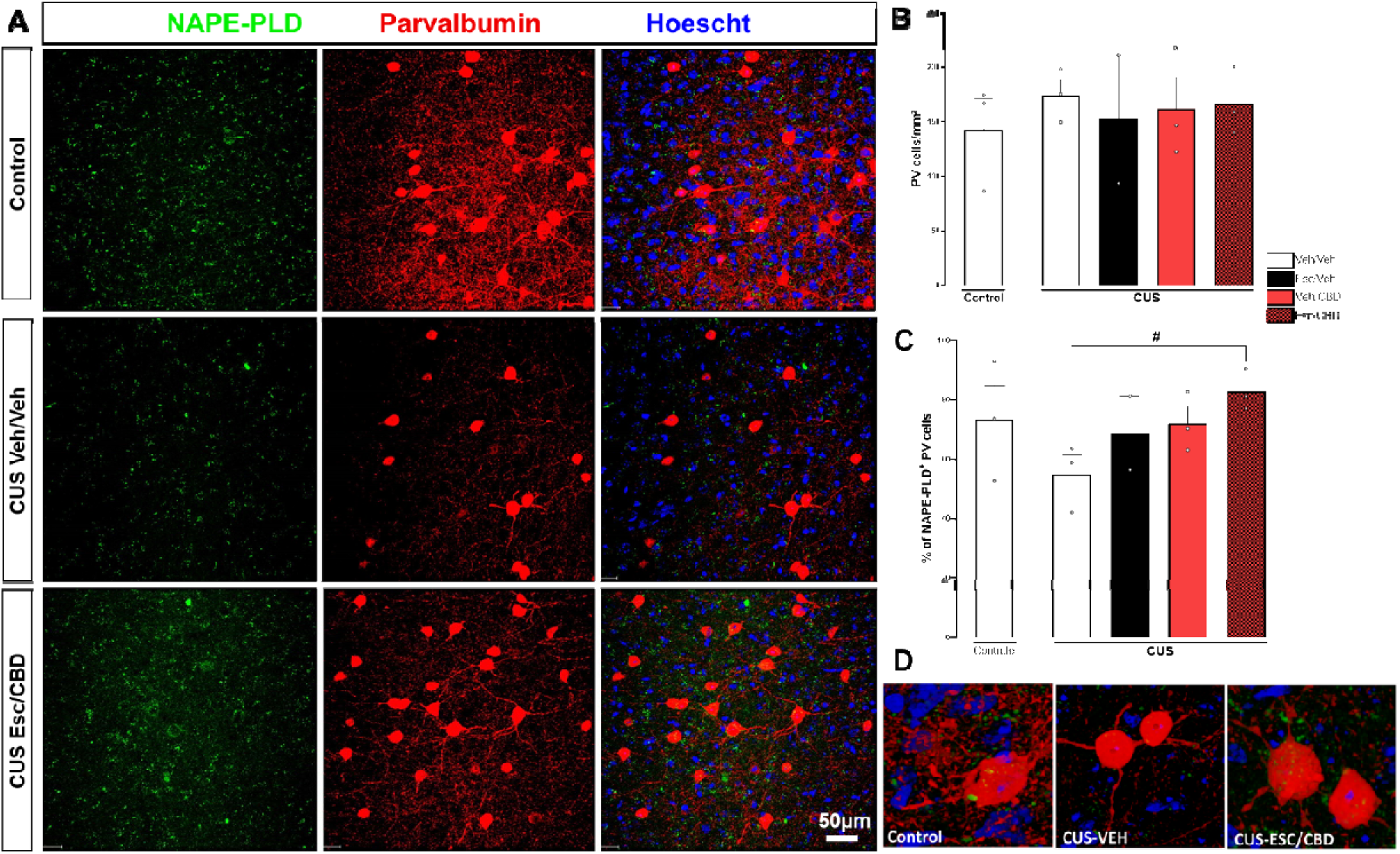
Escitalopram and CBD increase the density of NAPE-PLD-expressing parvalbumin (PV) interneurons in the PFC of stressed mice. (A) Representative images of NAPE-PLD and PV immunostaining in the PFC; (B) Density of PV-positive cells and (C) percentage of PV cells expressing NAPE-PLD in the PFC of stressed mice treated for 7 days with the combinations of vehicle or escitalopram (10mg/kg) with vehicle or CBD (7.5mg/kg). (D) Representative images of changes in PV+ cells expressing NAPE-PLD close to the soma induced by CUS and restored by the ESC+CBD treatment. Data represented as the Mean ± SEM; (#) indicates p<0.05 in comparison to the CUS Veh/Veh group (Two-way ANOVA; One-way ANOVA followed by Duncan; t test for independent samples).

**Figure 5.**
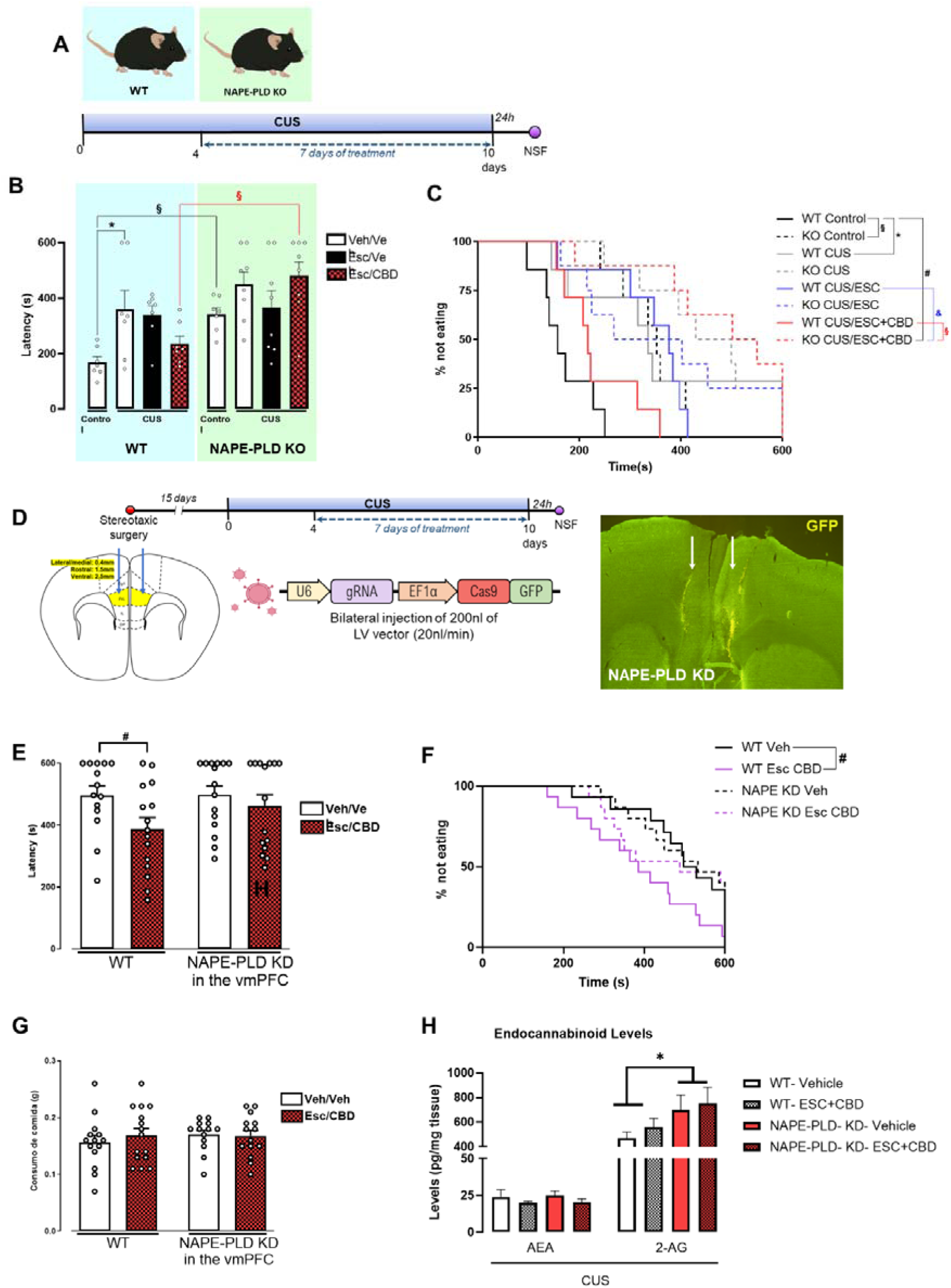
The anxiolytic-like effect of escitalopram and CBD is absent after genetic manipulations in NAPE-PLD expression. (A) Schematic representation of the timeline of the experiment involving NAPE-PLD knock out (K.O). mice. (B) Latency to feed in the novel environment of Wild type (W.T.) and NAPE-PLD and (C) percentage of mice not eating throughout the 600s of the NSF test in WT and NAPE-PLD KO mice exposed to 10 days of stress and 7 days of treatment with vehicle or escitalopram (10mg/kg) combined or not with CBD (7.5mg/kg); (D) Schematic representation of the viral vectors directing the CRISPR-Cas9-mediated deletion of NAPE-PLD and injection site of the viral vectors evidenced through the detection of GFP-positive cells; (E and F) Latency to feed in the novel environment, percentage of WT or NAPE-PLD knock down (K.D) mice not eating throughout the 600s of the NSF test and (G) food consumption in the home-cage of mice after 10 days of CUS and 7 days of treatment with Vehicle or with the combination of Escitalopram and CBD in which NAPE-PLD was knocked-down (K.D.) in the PFC (H) levels of the endocannabinoids, Anandamide (AEA) and 2-AG in the PFC of W.T. or NAPE-PLD K.D. mice. Data represented as Mean ± SEM; (*) indicates p<0.05 in comparison to the Control group corresponding to the same genotype; (§) indicates p<0.05 in comparison to the WT group corresponding to the same condition and treatment; (&) indicates p<0.05 in comparison to the CUS group treated only with escitalopram; (#) indicates p<0.05 compared to the stressed group treated with vehicle corresponding to the same genotype (Two-way ANOVA; One-way ANOVA followed by Duncan; Mantel-Cox log-rank test followed by Gehan-Breslow-Wilcoxon test).

### 3.4. Disruption of NAPE-PLD function impairs the anti-stress effects of escitalopram combined with cannabidiol: involvement of prefrontal cortex

Continuing the investigation of the mechanisms associated with the behavioral effects of ESC+CBD and based on results present in figures 4 and 5, we hypothesized that NAPE-PLD expression is required for the synergism of CBD on ESC anti-stress effects. Baseline latency to feed was higher in the NAPE-PLD KO mice compared to WT control mice (Figure 5B-C). Next, we submitted NAPE-PLD KO mice or WT control mice to a 10-day protocol of CUS (Figure 6A). In WT mice, but not in NAPE-PLD KO mice, CUS exposure significantly increased the latency to feed in the new environment in the NSFT (independent samples t test; WT Control vs WT CUS: t_7.04_=2.672, p=0.032; NAPE-PLD KO Control vs NAPE-PLD CUS: t_13_=2.032, p=0.063). The WT mice response after CUS in the NSF test was significantly different from the NAPE-PLD KO mice’s response (t_12_=5.589, p<0.001) (Figure 5 A-C).

Considering only the stressed groups, Two-way ANOVA indicated a significant effect of the factor genotype (F_1,39_=8.519, p=0.006), but no effect of Treatment (F_2,39_=0.644, p=0.531) and there was no significant interaction between factors (F_2,39_=2.475, p=0.097 Duncan posttest indicated that the WT CUS group treated with Escitalopram + CBD was significantly different from the NAPE-PLD CUS group that received the same treatment combination (F_5,39_=2.947, p=0.024) (Figure 5A-B). Consistent with these results, Mantel-Cox log-rank test (p<0.0001) followed by Gehan-Breslow-Wilcoxon test revealed that the response of the WT Control group was significantly different from the WT CUS group (p=0.013) and from the NAPE-PLD KO Control group (p<0.001). Interestingly, the response of the WT CUS group treated only with Escitalopram was significantly different from the WT CUS Escitalopram + CBD group (p=0.041). The response of WT CUS Escitalopram + CBD group was significantly different from the NAPE-PLD KO CUS Escitalopram + CBD group (p=0.003) (Figure 5C). Taken together, these results show that the combination of escitalopram and CBD fails to promote an anxiolytic-like effect in stressed mice with constitutive KO of NAPE-PLD.

We next wondered if the NAPE-PLD signaling within the PFC could explain the absence of the effects of the combination of ESC+CBD observed in NAPE-PLD K.O mice. Therefore, we tested our stress protocol and treatment regime in mice that received the intra-PFC injection of a viral vector directing a CRISPR/Cas9-construct-induced Knock-down (KD) of the NAPE-PLD gene in the PFC (Figure 5D, supplementary figure 5). In NAPE-PLD-PFC-KD mice, two-way ANOVA indicated that there was no effect of the factor genotype (F_1,54_=2.946, p=0.095), but there was a significant effect of the factor treatment (F_1,54_=8.075, p=0.008). The combination of escitalopram and CBD decreased the latency to eat in a novel environment in WT animals (t_27_=2.231, p=0.034), but the effect of escitalopram+CBD disappeared in NAPE-PLD vmPFC KD mice (t_28_=0.781, p=0.4412) (Figure 5E). Additionally, Mantel-Cox test followed by Gehan-Breslow-Wilcoxon test indicated a significant difference in the response of the WT group treated with the combination of escitalopram + CBD in comparison to the WT Veh-treated group in the NSF test (Mantel-Cox: p=0.050; Gehan-Breslow-Wincoxon: p=0.025) (Figure 5F). No differences were observed on food consumption post-NSF test (Figure 5G). The results suggest that the anti-stress effects of the combination of CBD plus escitalopram are absent after disruption of NAPE-PLD expression specifically in the PFC.

In addition to the behavioral analysis, we measured content of the AEA and 2-AG in the PFC (globally, not restricted to the site of viral vectors infusions) of NAPE-PLD-PFC-KD mice (Figure 5H). However, we did not observe any changes in AEA levels in NAPE-PLD mice. Interestingly, regarding 2-AG, we found a significant increase in its levels in the PFC of NAPE-PLD-KD mice (genotype: F(1,7) = 7.45, p < 0.05), independent of treatment status (treatment: F(1,7) = 3, p = n.s.) or genotype × treatment interaction (F(1,7) = 1.1, p=n.s). These findings suggest a compensatory increase in 2-AG production triggered by the local disruption of NAPE-PLD function in stressed mice and are consistent with previous results in another brain region, the ventral hippocampus, where NAPE-PLD deletion in stressed-trapped neurons also led to elevated 2-AG levels (Tevosian et al., 2023).

We also investigated the effect of DAGLα deletion in a similar protocol. Conversely, the CRISPR/Cas9-mediated deletion of DAGLα in the vmPFC did not prevent the anti-stress effects of Escitalopram + CBD (F_1,34_=8.075, p=0.008, Sup Figure 5). Consistent with this result, Mantel-Cox log-rank test followed by Gehan-Breslow-Wilcoxon indicated a significant difference in the responses of the DAGLα KD Veh-treated and the DAGLα KD Escitalopram + CBD-treated groups in the NSF test (Mantel-Cox: p=0.001; Gehan-Breslow-Wilcoxon: p=0.011) (Sup Figure 5). No differences were observed in the food-consumption in the home-cage after DAGLα PFC KD (Sup Figure4). Determination of 2-AG and AEA levels in the whole PFC of DAGLα PFC KD, did not show any differences considering the genotype, treatments or their interaction (Two-way ANOVA) (Sup. Table 2).

## 4. Discussion

The results presented in our study introduce the possibility of combining a low dose of CBD with escitalopram to enhance escitalopram’s efficacy by reducing its onset of action from 21 to 7 days in the context of heterotypic and homotypic protocols of stress. CBD (30mg/kg, i.p) induced an anti-stress effect in a 7 day-protocol of CUS while the behavioral effects of the SSRI escitalopram were only observed after 14 days at a higher dose (20mg/kg) or 21 day at a lower dose-10mg/Kg (Ribeiro et al., 2023). Previous studies using chronic stress models had shown anti-stress effects of CBD emerging after 14-(Campos et al., 2013. Fogaça et al., 2018) or 28-days treatment (Gáll et al., 2020) in rodents. In addition, Xu and colleagues demonstrated that 4 administrations of CBD (given weekly from the second week of CUS) induce an antidepressant-like effect in the forced swim test (FST) (Xu et al., 2020). It has been proposed that CBD could be a fast-acting antidepressant-like agent, like ketamine, using the forced swimming test (FST) as a readout as well as in the olfactory bulbectomy and learned helplessness animal models (Linge et al., 2016, Sales et al., 2019). However, this potential fast-acting property has been poorly investigated in models of chronic stress. Chronic stress is a major environmental factor and recognized as one of the leading risk factors for the development of anxiety and major depressive disorders (Mundt et al., 2000), for which antidepressants (ADs) are the first-line treatment (Cipriani et al., 2018). Given the well-documented delayed onset of therapeutic response to SSRIs in these conditions, our findings may have important clinical implications.

When SSRI monotherapy fails, a commonly adopted strategy in psychiatric patients is combination therapy with augmentation drugs, such as other antidepressants, antipsychotics, or mood stabilizers (Moret et al., 2005, Palaniyappan et al., 2009, Nuñez et al., 2022). Our study is the first to describe a potential benefit of the combination of escitalopram with CBD in models of repeated stress. Drug combinations often allow reduced drug doses, potentially decreasing the incidence and/or severity of side effects, which should improve patient adherence and outcomes. Here, we find that adding one-quarter of CBD’s effective dose in CUS protocols (Campos et al., 2013; Fogaça et al., 2018) enhances the anti-stress efficacy of a low dose of escitalopram (10 mg/kg), which, when administered alone, only produces observable effects after 21 days in the chronic stress protocol (Ribeiro et al., 2023). In addition, the fact that our combination treatment regimen induced an anxiolytic-like effect in several different models of stress reinforces the robustness of the effect, especially considering that heterotypic and homotypic stress protocols involve partially different neurobiological mechanisms and differ on their neuroendocrine outcomes (Morena et al., 2016; Kopp et al., 2013; Scarante et al., 2017, Sabban & Serova, 2007).

Putative interactions between CBD and conventional antidepressants have also been reported with other cannabinoid-related compounds, but these studies used acute treatment regimens (with higher doses) and relied on the FST as a single behavioral readout [39–41]. Accordingly, using a behavioral test based on a similar behavioral paradigm, the TST, we demonstrated that mice exposed to the SDS, the combination of ESC+CBD reduces the latency for the first immobility episode. However, it is important to emphasize that the FST and TST measure acute treatment responses, which differ from the clinical scenario where 4-12 weeks of treatment are needed. In the last decade several authors have discussed that the immobility time measured during the FST and TST are more indicative of adaptive behaviors (passive vs. active coping in response to acute stress) rather than reflecting a chronic state of depressive-like behaviors (Molendijk et al., 2015). This suggests caution when interpreting the efficacy of drugs in FST/TST whose therapeutic effects typically require chronic treatment to manifest in humans. Notably, novelty-suppressed feeding, the main behavioral readout used in our study, uniquely requires chronic, rather than acute, antidepressant treatment to reverse CUS-induced impairments, thereby reflecting the delayed therapeutic response observed in clinical practice (Samuels and Hen, 2011).

Previous work of our group suggested that 14-days of treatment with CBD (30mg/Kg) increases hippocampal AEA levels (Campos et al., 2013; Leweke et al., 2012) a mechanism also observed after the administration of a FAAH inhibitor, such as URB597. URB597 induced an anxiolytic-like effect in mice exposed to SDS (Fusse et al., 2024). However, our results indicated that that, while the combination of escitalopram with a subeffective dose of CBD (7.5mg/Kg) induced an anxiolytic-like effect after 7 days of treatment, the same was not seen after the combination of escitalopram with a subeffective dose of URB597 (dose of 0.1mg). This may be related to differences in the pharmacological potency of these drugs in affecting the endocannabinoid system associated with the multitarget characteristic of CBD (Campos et al., 2012). Antidepressants can also modulate endocannabinoid levels, such as AEA (Hill et al., 2006), and their effects are potentiated in FAAH-deficient mice (Bambico et al., 2010), it is reasonable to hypothesize that the add-on effects of CBD in accelerating escitalopram’s anti-stress response might involve brains endocannabinoid signaling.

Leishman and colleagues (2018) compared the molecular effects of URB597 and CBD using lipidomic analysis and concluded that they differentially modulated the levels of NAPE-PLD (the main enzyme responsible for producing AEA in the adult brain) products, suggesting that CBD and URB597 (i.e., FAAH inhibition) may have distinct impacts on anandamide levels, with the former increasing anandamide synthesis and the latter decreasing its degradation. Our results suggested that the treatment with the combination of ESC+CBD seems to counteract the negative effect of stress on NAPE-PLD expression in the PFC, suggesting the participation of NAPE-PLD in the behavioral outcomes of the combination of escitalopram and CBD. We observed that the combination of escitalopram and CBD did not promote an anxiolytic-like effect in NAPE-PLD KO mice or in KD mice in which the expression of NAPE-PLD was reduced in the PFC via CRISPR-Cas9. Interestingly, both escitalopram and CBD have been shown to alter the activity and expression of NAPE-PLD (Leishman et al., 2018; Smaga et al., 2019).

Our data indicated that stress decreases the expression of NAPE-PLD in the PFC and that the combination of ESC+CBD counteracts this effect. The relevance of NAPE-PLD for stress responses has been previously investigated. Chen and colleagues (2023) demonstrated that NAPE-PLD KO mice showed a depressive-like phenotype in the sucrose preference test and we found increased latency to feed in the NSFT test (Figure 6B). Tevosian and colleagues (2023) deleted NAPE-PLD expression only in neurons activated during exposure to SDS. In this context, NAPE-PLD conditional KD exhibited an anxiogenic-like phenotype.

Unexpectedly, chronically stressed mice with NAPE-PLD knockdown in the PFC exhibited elevated levels of 2-AG in this region. Accordingly, a previous study where NAPE-PLD was only downregulated in neurons recruited during stress responses, also observed an increase in 2-AG levels in the ventral portions of the hippocampus. Acute and homotypic chronic stressors can increase 2-AG levels in the prefrontal cortex and amygdala, likely as a consequence of stress-induced glucocorticoid activity (Bedse et al., 2020; Patel et al., 2009; Scarante et al., 2017). Our findings suggest that impaired NAPE-PLD activity disrupts endocannabinoid signalling under chronic stress, leading to a compensatory increase in 2-AG. This response may serve to counteract glucocorticoid actions and the enhanced inhibitory tone observed in PFC pyramidal neurons during the stress response (Ferguson and Gao, 2018; Nawreen et al., 2024), a mechanism considered essential for the adaptation of neuroendocrine responses to stress (Hill et al., 2010).

Indeed, we observed that chronic stress reduced dendritic spine density in cortical pyramidal neurons, supporting the notion that our protocol induced some degree of prefrontal cortex (PFC) hypofunction. Shepard and Coutellier (2018) reported that two weeks of CUS decreased glutamatergic activity in the PFC, accompanied by enhanced NMDA receptor activity, specifically in parvalbumin-positive interneurons. This increase in local inhibition mediated by PV interneurons can impair the synaptic plasticity of pyramidal neurons and may account for the reduced number of dendritic spines observed in stressed mice.

Interestingly, we found that the combination of escitalopram and CBD seems to restore NAPE-PLD expression after CUS, especially in PFC PV+ interneurons. PV + interneurons are major regulators to maintain appropriate excitatory/inhibitory balance in cortex (Ferguson and Gao, 2018). Page and colleagues (2019) showed that the exposure to 4 weeks of CUS significantly increased the activity of PV-positive cells in the PFC. They also demonstrated that the chemogenetic activation of PV-positive cells for 21 days in the PFC was sufficient to promote an anxiogenic-like phenotype in female mice (Page et al., 2019), while the chemogenetic inhibition of this population of interneurons mitigates the behavioral effects of 14-day protocol of chronic stress (Fogaça et al., 2021; Nawreen et al., 2020, 2024). Antidepressant treatment modulates the dendritic branching and the activation pattern of PV-positive interneurons (Gilabert et al., 2013; 2017). Moreover, the fast-acting antidepressant-like effect of ketamine is associated with an increased activation of pyramidal neurons via the inhibition of fast-spiking interneurons in the PFC (Zhang et al., 2021).

In line with the findings of Kucera et al. (2018), we observed that NAPE-PLD is expressed closely in the soma of PV-positive interneurons. Considering that cortical PV+ interneurons receive excitatory glutamatergic inputs directly onto their somata (Keller & White, 1987; Milicevic et al., 2024), a stress-induced reduction in NAPE-PLD expression could compromise this self-regulatory inhibitory mechanism, thereby leading to increased PV+ interneuron activity within the PFC. This suggests that somatic recruitment of NAPE-PLD-mediated endocannabinoid synthesis may play a critical role in modulating glutamatergic drive onto PV interneuron cell bodies. Alternatively, AEA synthesized in PV+ neurons might act in a paracrine manner on neighboring cells (Bacci et al., 2004). Moreover, PV+ interneurons themselves could provide negative feedback through GABA release onto pyramidal cell somata, thereby dampening their own activity and contributing to the restoration of excitatory-inhibitory balance in the PFC.

Thus, we propose that CBD enhances the behavioral effects of ESC by restoring somatic NAPE-PLD expression of PV+ cells of the PFC. This restoration reestablishes endocannabinoid tone in the region, promoting the self-inhibition of PV interneurons and consequently correcting the excitatory–inhibitory imbalance. The resulting disinhibition of pyramidal neurons would favor neuroplasticity, as evidenced by the recovery of synaptic markers such as synaptophysin and the normalization of dendritic spine density, ultimately leading to improvements in anxiogenic- and depressive-like behaviors. Moreover, the combined effects of ESC and CBD may also engage alternative metabolic pathways of unsaturated fatty acids, resulting in the formation of NAEs and NAANs via NAPE-PLD activity (Leishman et al., 2018).

Finally, although not directly explored in this study, we cannot rule out the contribution of serotonergic mechanisms to the behavioral effects of CBD + ESC in PV-positive neurons. This possibility is particularly relevant given the established role of 5-HT1A receptor activation in the behavioral effects of both SSRIs and CBD (Campos et al., 2008, 2012; Fogaça et al., 2013) and in mediating plasticity within the prefrontal cortex (Higa et al., 2022). In the PFC, 5-HT1A receptors are abundantly expressed at the axon initial segment of pyramidal neurons and in PV+ interneurons (De Felipe et al., 2001). Similar to cannabinoid receptors, 5-HT1A receptors are coupled to Gi/o proteins; their activation in pyramidal neurons suppresses action potential generation, contributing to PFC hypofunction as observed in chronically stressed mice (De Felipe et al., 2001). Moreover, Fogaça et al. (2018) demonstrated that the 5-HT1A antagonist WAY-100635 fails to block the behavioral effects of CBD in stressed mice, while cannabinoid receptors were implicated in mediating these anti-stress actions.

As with any study, our findings have some limitations. In the control non-stressed groups (SDS and CUS) of our study, two response patterns emerged: most animals responded within 400 seconds, while a subgroup reached the 600-second cut-off, a profile typically seen in stressed mice. This variability may reflect procedural stress (daily injections), environmental factors, or individual coping differences. The lack of clear stress effects in behavioral tests is a common but still underreported challenge in preclinical research. To avoid adding further stressors, we did not isolate animals after procedures or expose them to repeated daily stress. Despite these factors, the ESC–CBD combination consistently produced a robust anxiolytic-like effect, with almost all treated mice showing latencies below 300 seconds. Another limitation is the absence of a non-stressed group in the conditional KD protocol; for technical and ethical reasons, we focused on the role of NAPE-PLD in treatment effects. Finally, we did not directly measure NAPE-PLD activity or fully dissect the pharmacological mechanisms by which escitalopram and CBD modulate their expression. Future studies should address these issues, expand dose-response analyses, and test generalizability of our finding across mice of sex and age.

## Conclusion

Altogether, our pre-clinical data indicate that CBD could be a viable strategy as an add-on therapy to optimize the therapeutic response to antidepressants. We propose that CBD could accelerate escitalopram’s anti-stress effects via the recruitment of NAPE-PLD -mediated signaling by restoring the inhibitory/excitatory tons via self-inhibition of PV+ neurons located within the prefrontal cortex (Figure 7).

**Figure 7.**
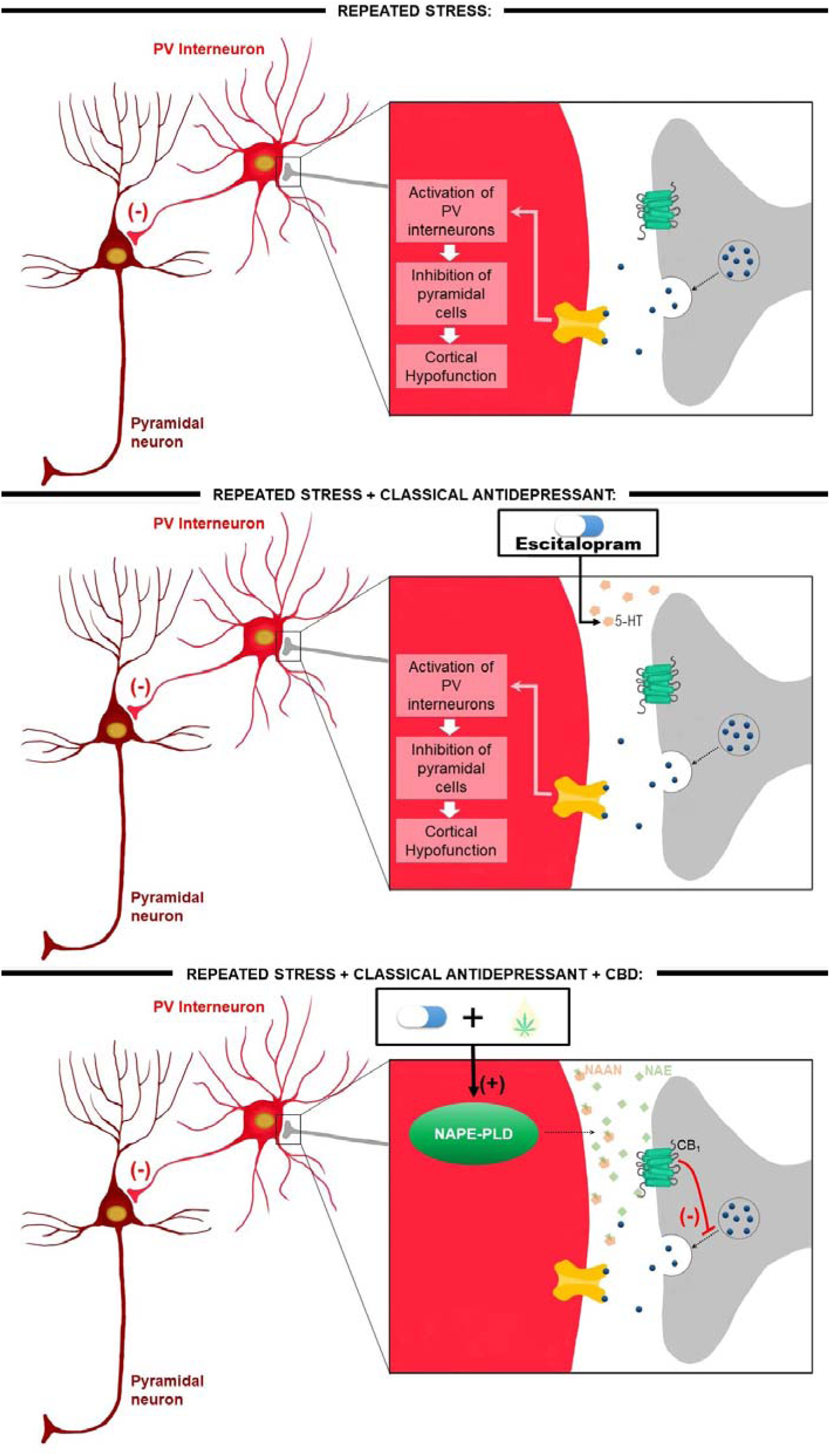
Proposed neurochemical mechanisms underlying the anti-stress effects of escitalopram combined with sub-effective doses of cannabidiol. In the stressed brain, hyperactivation of PV interneurons disrupts the excitatory–inhibitory balance in the prefrontal cortex (PFC), leading to reduced pyramidal neuron activity. Short-term treatment with classical antidepressants such as escitalopram alone is insufficient to reverse this stress-induced cortical inhibition. In contrast, escitalopram combined with CBD may recruit NAPE-PLD activity, restoring endocannabinoid tone (AEA, 2-AG, N-acylethanolamines [NAEs], and N-acyl amino acid neurotransmitters [NAANs]). These bioactive lipids act on multiple targets, including presynaptic cannabinoid receptors on excitatory terminals projecting to PV interneurons. Their activation reduces glutamate release, thereby dampening PV interneuron activity, relieving pyramidal neurons from excessive inhibition, and ultimately restoring synaptic plasticity in the PFC.

## Acknowledgments

We would like to thank Eleni Tamburus, Giuliana Bertozi and Tadeu Franco Vieira (*in memoriam*) for their excellent technical support. We want to express our gratitude to our lab colleagues for keeping an amazing and collaborative environment. We give special thanks to Prof. Serge Luquet for providing the NAPE-PLD K.O. mice. This study was supported by the following Brazilian public research funding agencies: the São Paulo Research Foundation (FAPESP), the Brazilian National Council for Scientific and Technological Development and the Coordination for the Improvement of Higher Education Personnel (CAPES). The research funding was originated from the FAPESP Young Investigator Grant (2015/05551-0), FAPESP Thematic Grant (2017/24304-0), the L’Oreal-UNESCO Brazilian Academy of Science for Women in Science Fellowship, and a CNPq Universal Grant line 2 and 1D (400033/2016-0). FFS and ACC received a FAPESP fellowship (2019/09178-3; 2021/13515-5, 2022/13764-8). ACC and FSG are recipients of CNPq individual research fellowships (304336/2022-0).

## Authorś Contribution: CRediT

**Franciele Franco Scarante:** Conceptualization, Methodology, Validation, Formal Analysis, Investigation, Writing – Original draft, Writing – Review and editing, Visualization. **Vinícius Detoni Lopes:** Conceptualization, Methodology, Validation, Formal Analysis, Investigation, Writing – Original draft. **Eduardo J. Fusse:** Methodology, Investigation. **Maria Adrielle Vicente:** Methodology, Investigation, Writing – Original draft. **Gabriel H. D. de Abreu**: Investigation. **Pedro H. C Lirio**: Investigation**. Viviani Nardini:** Methodology, Investigation. **Carlos Arterio Sorgi:** Methodology, Investigation. **Lucia H. Faccioli:** Methodology, Investigation, Writing – Review and editing. **Katiuchia U Sales:** Investigation, Writing – Review and editing. **Jaime E.C. Hallak:** Writing – Original draft, Writing – Review and editing. **Jose A Crippa:** Writing – Original draft, Writing – Review and editing **Feng Gou:** Conceptualization, Writing – Original draft, Writing – Review and editing. **Hui-Chen Lu**-Conceptualization, Writing – Original draft, Writing – Review and editing. **Francisco S. Guimarães:** Conceptualization, Resources, Writing – Review and editing. **Kenneth Mackie:** Conceptualization, Writing – Original draft, Writing – Review and editing. **Alline C. Campos:** Conceptualization, Methodology, Formal Analysis, Resources, Writing – Original draft, Writing – Review and editing, Visualization, Supervision, Project administration, Funding acquisition.

## Data Statement

All data generated in this study is available upon formal request to the corresponding author.

## Conflict of interest

ACC, FSG, JECH and JAC are coinventors of the patent “Cannabinoid-containing oral pharmaceutical composition, method for preparing and using same,” INPI on September 16, 2016 (BR 112018005423-2). FSG, JECH and JAC are inventors of the patent “Fluorinated CBD compounds, compositions and uses thereof. Pub. No.: WO/2014/108899. International Application No.: PCT/IL2014/050023,” Def. US number Reg. 62193296; July 29, 2015; INPI on August 19, 2015 (BR1120150164927; Mechoulam R, Zuardi AW, Kapczinski F, Hallak JEC, Guimarães FS, Crippa JAS, Breuer A). Universidade de São Paulo (USP) has licensed this patent to Phytecs Pharm (USP Resolution No. 15.1.130002.1.1) and has an agreement with Prati-Donaduzzi to “develop a pharmaceutical product containing synthetic CBD and prove its safety and therapeutic efficacy in the treatment of epilepsy, schizophrenia, Parkinson’s disease, and anxiety disorders.” JAC is a member of the International Advisory Board of the Australian Centre for Cannabinoid Clinical and Research Excellence (ACRE) – National Health and Medical Research Council (NHMRC). JAC and JEH have received travel support to attend scientific meetings and personal consultation fees from BSPG Pharm. JAC and JECH have received personal consultation fees from BSPG-Pharm, and PurMed Global in the past. JAC received speaking fees from Torrent, Green Care Store, Janssen and consultation and speaking fees from EaseLabs. ACDC received speaking fee from EaseLabs. The other authors declare no competing interests.

## Notes

### Summary of Updates

In this revised version of the manuscript, we have removed the patient data and incorporated additional mechanistic and genetic experiments demonstrating the involvement of NAPE-PLD and PV neurons in mediating the effects of a subeffective dose of CBD, which acts synergistically with escitalopram to accelerate the onset of behavioral responses in chronically stressed male mice.

