## Supplemental material for "Cannabidiol reduces the latency for the behavioral effect of escitalopram in chronically stressed male mice: involvement of NAPE-PLD expressed in parvalbumin-positive interneurons and the prefrontal cortex"

**Methods**

*Tail suspension test (TST)*

The TST, a classical test to evaluate both passive and active coping behavior, employed in the present study was a modified version of that validated for NMRI mice (Steru et al., 1985). At the day of the test (10^th^ day of the stress protocol- SDS or CUS), all mice were carefully moved for short distance from the holding facility to the testing room and left there undisturbed for at least 3 h. Animals were individually suspended by the tail to a horizontal ring-stand bar (height from floor = 35 cm) using adhesive tape (distance from tip of tail = 2 cm) for 6 minutes. The test session was recorded, and the total immobility time was evaluated by an experimenter blinded to the treatments or condition (stressed or non-stressed). Mice often demonstrated several escape-oriented behaviors interspersed with bouts of immobility of increasing duration as the test session progressed.

*Doublecortin (DCX) detection and analysis*

For the DCX detection procedures, slices containing dorsal parts of the hippocampus of each experimental animal underwent to an antigen retrieval step in citrate buffer (10mM Citric Acid, 0.05% Tween 20, pH 6.0; at 70° C for 30 minutes) after receiving 3 washes in TBS. Slices were left for cooling down at room temperature and then incubated in a blocking solution (BSA 1% + 0.25% Triton 100X in TBS) for 2 h, at least. Afterwards, the slices were incubated overnight with the primary antibody (goat anti-DCX- Santa Cruz Biotechnology, 1:200) and subsequently incubated with the secondary antibody for 1h (1:1,000 Vectastain anti-goat biotinylated). We also performed an additional step of incubation with A+B complex for 1 h (1:1,000ABC Elite-Vectastain kit, Vector Labs - Burlingame, CA- United States) followed by staining with 3,3'-Diaminobenzidina (DAB 0.2mg/mL; Sigma-Aldrich, Missouri, United States) for 10 minutes, approximately. Finally, slices containing hippocampi were mounted on glass slides/coverslips with Permount (DCX-Fisher Scientific, Loughborough, United Kingdom) as mounting media. DCX+ cells were counted in a 40x objective of a light microscope (Olympus BX60) by an experimenter blinded for the treatments and condition (non-stressed or stressed). A cell was considered positive for DCX only if the soma was stained. Cells were counted in subgranular or granular zone of the Dentate Gyrus (DG). The total number of cells were normalized to the DG area determined with 10x objective and then the number of DCX+ was estimated by calculating the total hippocampal volume as determined by the sum of the areas of the sections multiplied by the distances between them (series of hippocampal sections located between 1.3 mm and 2.5 mm posterior to bregma) (Campos et al., 2013, 2015).

**Results:**

Regarding the expression of DCX+ cells/mm² in the dentate gyrus after 14 days of the experimental protocol, a statistically significant difference was found for the condition factor (F(1,29) = 5.591, p < 0.05). This result indicates that stress reduced the number of DCX+ cells compared to control animals, regardless of the treatment factor. A one-way ANOVA followed by Duncan’s post hoc test did not reveal a statistically significant difference between stressed animals treated with vehicle and control animals that received the same treatment. The analysis further indicated that repeated treatment with CBD for 14 days in control animals significantly increased the number of DCX+ cells in the hippocampus of adult mice (Sup. Table 1 and Sup figure 1). In contrast, treatment with CBD (30 mg/kg) or Escitalopram (20 mg/kg) in stressed animals did not alter the number of DCX+ cells (Sup. Table 1 and Sup figure 1). Statistical analysis of DCX+ cells located in the granular layer of the dentate gyrus, which suggests a stage of migrating cells, showed no difference for any of the factors tested (condition or treatment), nor for their interaction.

**Sup Table 1. Number of doublecortin-positive cells in the dentate gyrus.**

| Group | 7 Days | 14 Days |
| --- | --- | --- |
| Control + Vehicle | 118.3 ± 9.08 | 94.5 ± 12.23 |
| Control + Cannabidiol | 129.5 ± 23.63 | 156.4 ± 18.78 * |
| Control + Escitalopram | 112.7 ± 6.60 | 103.9 ± 14.88 |
| Stress + Vehicle | 89.8 ± 8.65 * | 104.2 ± 20.81 |
| Stress + Cannabidiol | 102.9 ± 11.67 | 86.5 ± 24.77 |
| Stress + Escitalopram | 127.4 ± 19.11 | 52.3 ± 21.63 |

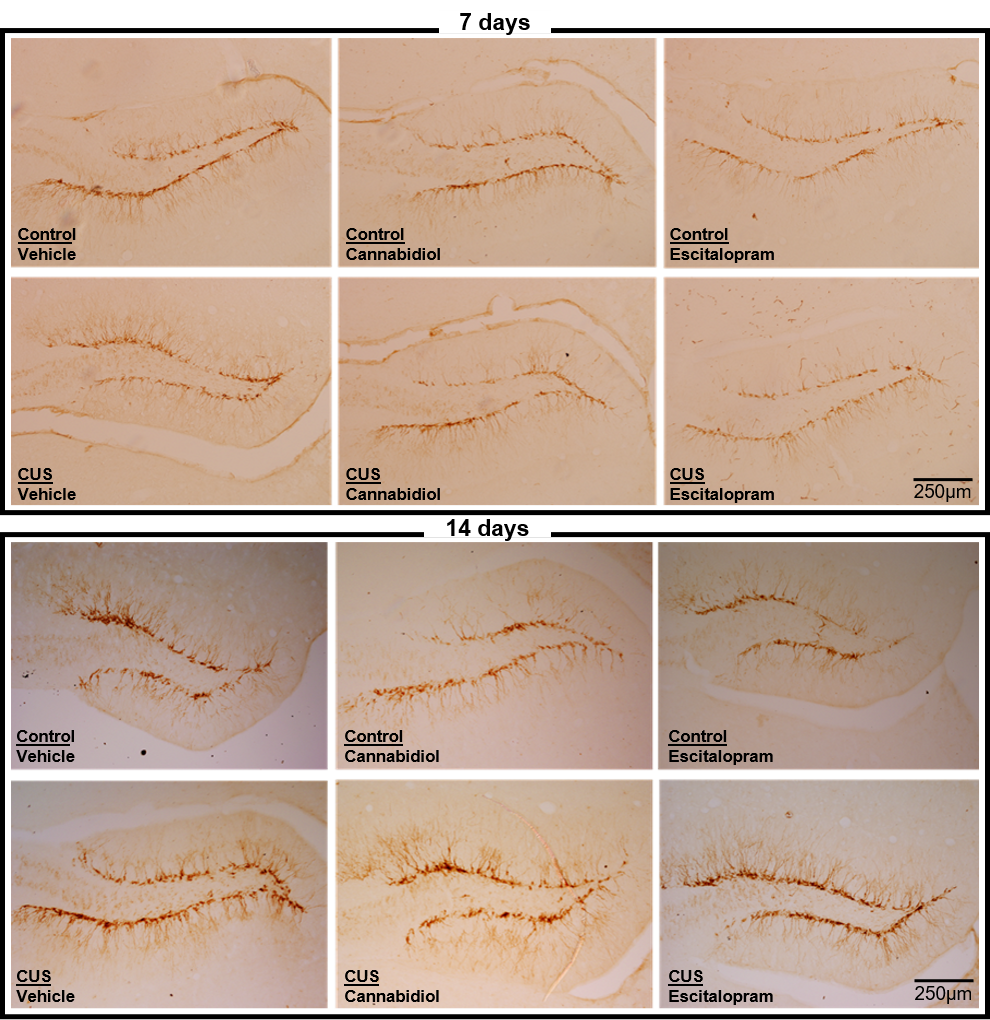

**Figure S1. Representative images of doublecortin-positive cells in the dentate gyrus of the hippocampus.** Immunolabeled DCX-positive cells in the dentate gyrus of mice submitted to 7 or 14 days of chronic unpredictable stress and treated with vehicle, escitalopram (20mg/kg) or cannabidiol (30mg/kg). Data represented as Mean ± SEM; (*) p<0.05 in comparison to the control group treated with vehicle (One-way ANOVA followed by Duncan).

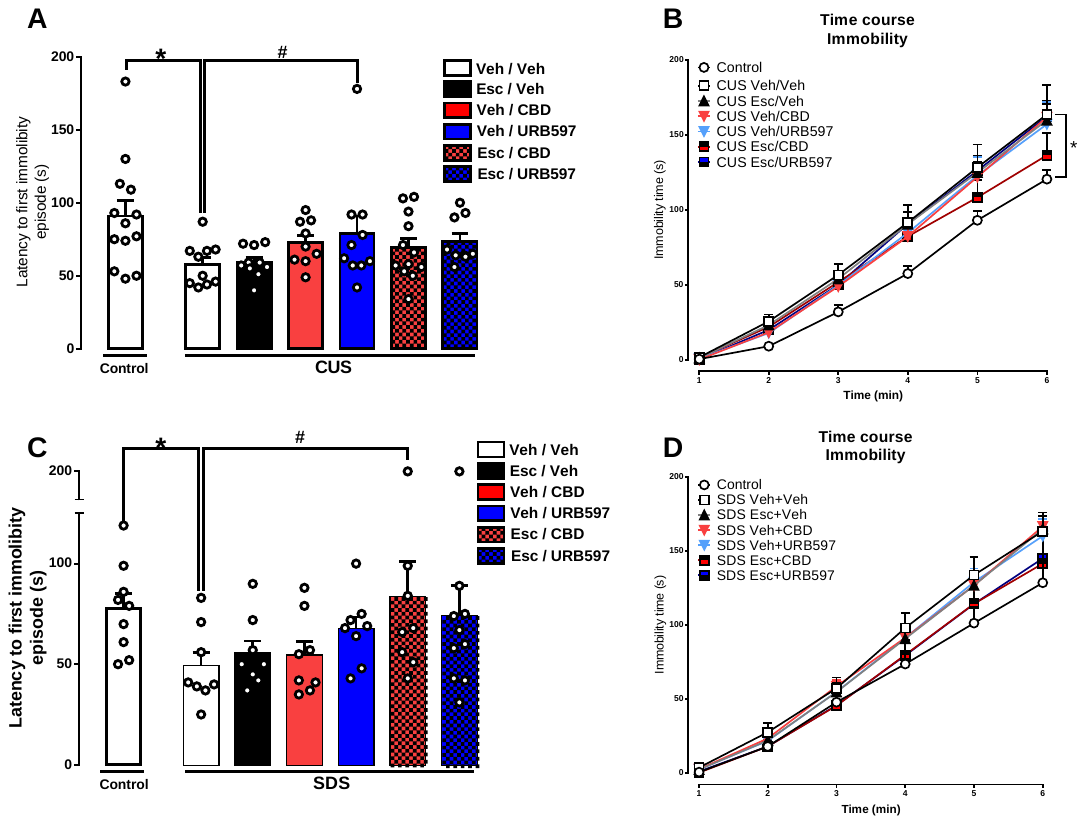

**Figure S2. Stress alters the coping strategies of mice in the TST.** Latency to the first immobility episode (A and C) and immobility time along the 6 minutes of the tail suspension test (B and D) in mice submitted to the chronic unpredictable stress (A and B) or to the social defeat stress (C and D) and treated with the combinations of vehicle or escitalopram (10mg/kg) with vehicle, CBD (7.5mg/kg) or URB597 (0.1mg/kg). Data represented as Mean ± SEM; (*) represents p<0.05 in comparison to the control (non-stressed) (t test for independent samples); (#) represents p<0.05 in comparison to the stressed group treated with vehicle.

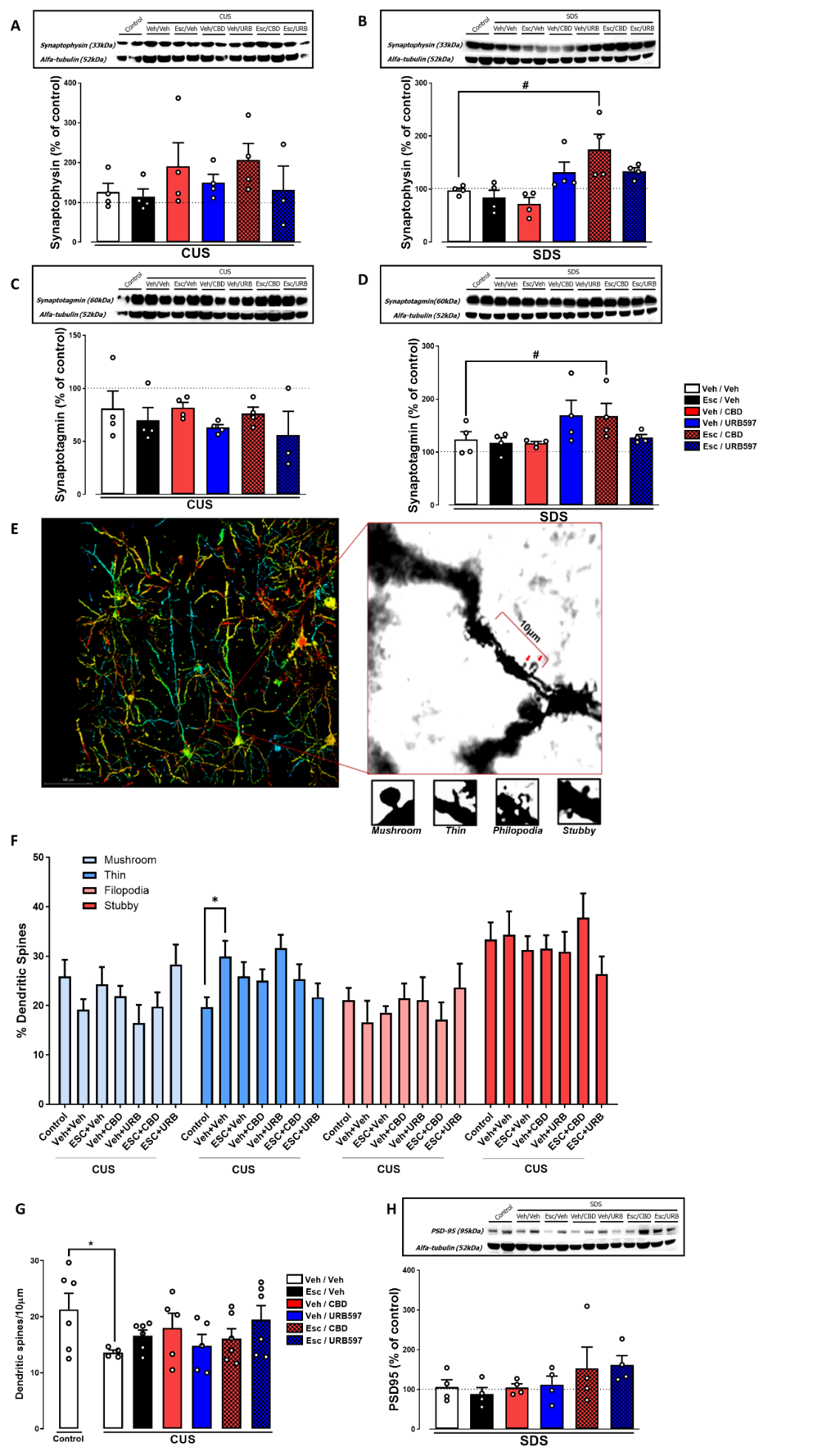

**Figure S3- Stress causes a postsynaptic impairment and combined CBD and escitalopram increases the expression of presynaptic proteins in the PFC.** Effect of the 7-day treatment with the different combinations of vehicle or escitalopram (10mg/kg) and vehicle, CBD (7.5mg/kg) or URB597 (0.1mg/kg) on the relative expression of the presynaptic proteins synaptophysin (A and B) and synaptotagmin (C and D) in mice submitted to stress. (E) Representative image of a pyramidal neuron in the PFC and explanation of the quantification and morphological analysis of dendritic spines; (F) Percentage of dendritic spines for each of the morphologies, relative to the total number of dendritic spines for each group; (G) Total number of dendritic spines in tertiary dendrites of mPFC pyramidal neurons in mice treated with the different drug combinations and submitted to stress; (H) Relative expression of the postsynaptic protein PDS95 in stressed mice treated with the different treatment combinations. Data represented as the Mean percentage relative to control (non-stressed group) ± SEM (A-D and H) or as Mean ± SEM (F-G); (*) indicates p<0.05 compared to the control group; (#) indicates p<0.05 compared to the stressed group treated with vehicle (Two-way ANOVA; One-way ANOVA followed by Duncan; t test for independent samples).

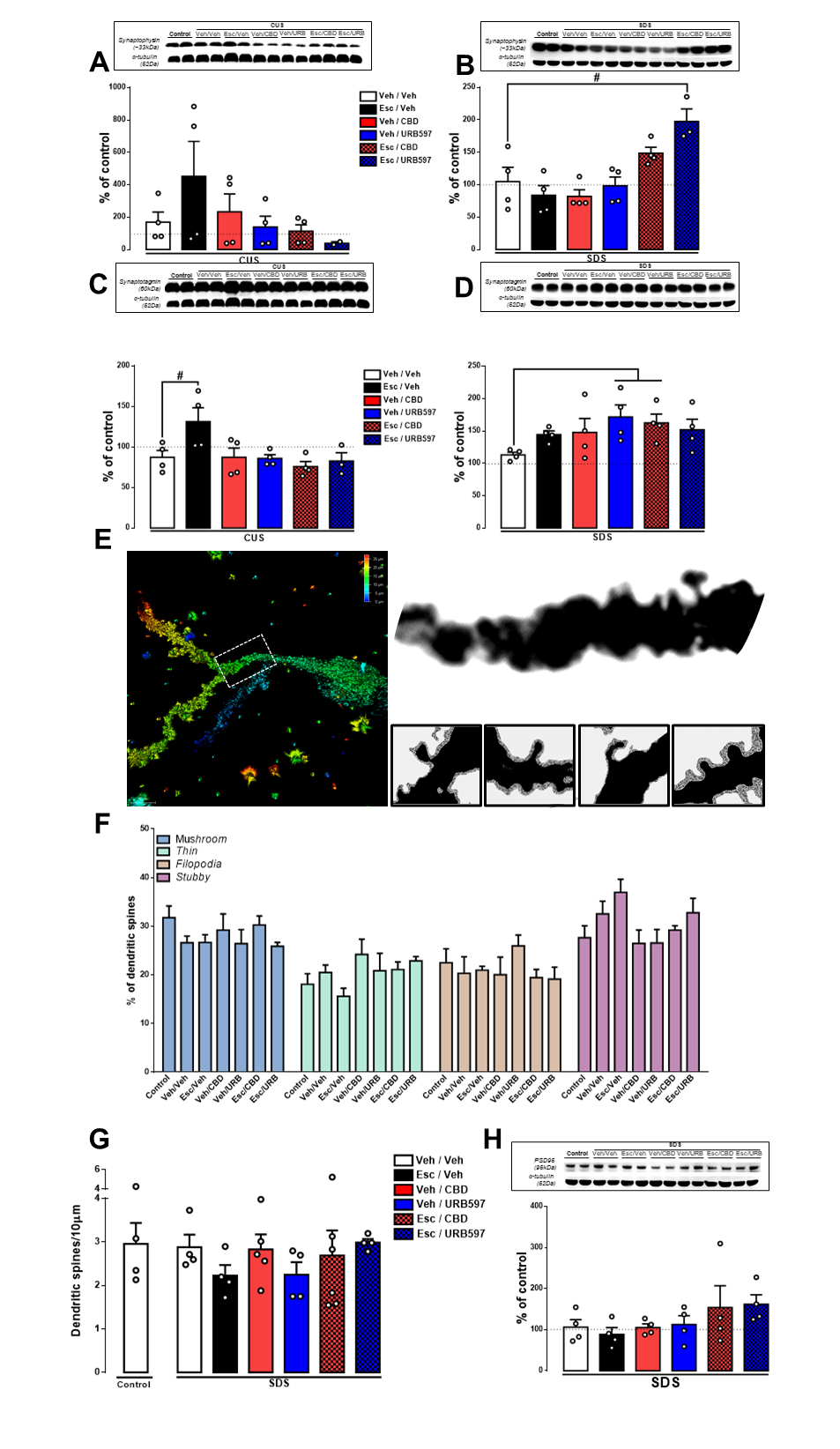

**Figure S4. Different treatment combinations induce alterations in hippocampal presynaptic proteins in stressed mice.** Relative hippocampal expression of the pre-synaptic proteins synaptophysin (A and B) and synaptotagmin (C and D); (E) Representative image of dentage gyrus granular neuron and classification of dendritic spines; (F) Percentage of dendritic spines per morphology and (G) total number of dendritic spines in secondary dendrites of granular neurons of the dentate gyrus of the hippocampus in mice; (H) relative hippocampal expression of PSD-95 in mice submitted to chronic unpredictable stress (CUS) (A and C) or social defeat stress (B, D, E-H) and treated with different combinations of vehicle or escitalopram (10mg/kg) or vehicle, cannabidiol (7.5mg/kg) or URB597 (0.1mg/kg). Data represented as the Mean percentage relative to control (non-stressed group) ± SEM (A-D and H) or as Mean ± SEM (F-G); (#) represents p<0.05 in comparison to stressed group treated with vehicle (Two-way ANOVA; One-way ANOVA followed by Duncan).

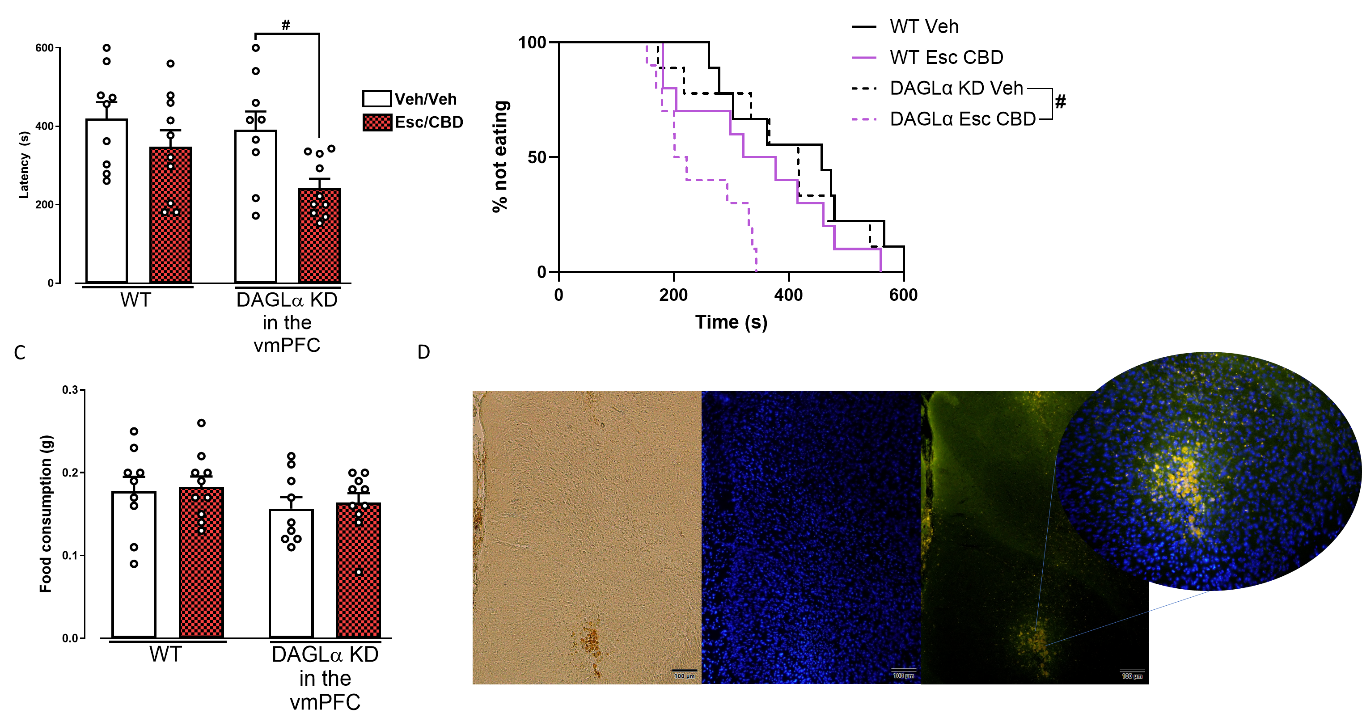

**Figure S5-** **DAGL is not involved in the behavioral effects of CBD+ESC treatment.** (A) Latency to feed in the novel environment, (B) percentage of mice not eating throughout the 600s of the NSF test and (C) food consumption in the home-cage of mice after 10 days of CUS and 7 days of treatment with Vehicle or with the combination of Escitalopram and CBD in which DAGL was (D) knocked-down (KD) in the PFC. Data represented as Mean ± SEM; (#) indicates p<0.05 compared to the stressed group treated with vehicle corresponding to the same genotype (Two-way ANOVA; One-way ANOVA followed by Duncan; Mantel-Cox log-rank test followed by Gehan-Breslow-Wincoxon test).
